# Rats strategically manage learning during perceptual decision making

**DOI:** 10.1101/2020.09.01.259911

**Authors:** Javier Masís, Travis Chapman, Juliana Y. Rhee, David D. Cox, Andrew M. Saxe

**Author notes:** Princeton Neuroscience Institute, Princeton University, Princeton, NJ 08544, U.S.A. MIT-IBM Watson AI Lab, Cambridge, MA 02142, U.S.A.

## Abstract

Balancing the speed and accuracy of decisions is crucial for survival, but how organisms manage this trade-off during learning is largely unknown. Here, we track this trade-off during perceptual learning in rats and simulated agents. At the start of learning, rats chose long reaction times that did not optimize instantaneous reward rate, but by the end of learning chose near-optimal reaction times. To understand this behavior, we analyzed learning dynamics in a recurrent neural network model of the task. The model reveals a fundamental trade-off between instantaneous reward rate and perceptual learning speed, putting the goals of learning quickly and accruing immediate reward in tension. We find that the rats’ strategy of long initial responses can dramatically expedite learning, yielding higher total reward over task engagement. Our results demonstrate that prioritizing learning can be advantageous from a total reward perspective, and suggest that rats engage in cognitive control of learning.

## Introduction

The speed-accuracy trade-off in decision making has been the subject of intense research, dating back nearly one-hundred years [1–14]. When facing noisy perceptual inputs, the longer an agent takes in making a choice the more likely that choice will be advantageous, but the less time is left to tackle subsequent choices. Choosing the right amount of time to deliberate on a particular decision is crucial for maximizing reward rate [9, 10].

Studies of the speed-accuracy trade-off have focused on how the brain may solve it [9, 15], what the optimal solution is [10], and whether agents can indeed manage it [11, 14, 16–22]. Though most work in this area has taken place in humans and non-human primates, several studies have established the presence of a speed-accuracy trade-off in rodents [23–28]. The broad conclusion of much of this literature is that after extensive training, many subjects come close to optimal performance [11, 17–21, 29–31]. When faced with deviations from optimality, several hypotheses have been proposed, including error avoidance, poor internal estimates of time, and a minimization of the cognitive cost associated with an optimal strategy [10, 30–32].

Past studies have shown how agents behave after reaching steady state performance[11, 17–21, 29, 30], but relatively less attention has been paid to how agents learn to approach near-optimal behavior (*but see* [18, 33]). While maximizing instantaneous reward rate is a sensible goal when the task is fully mastered, it is less clear that this objective is appropriate during learning.

Here, we set out to understand how agents manage the speed-accuracy trade-off during learning by studying the learning trajectory of rats in a free response two-alternative forced-choice visual object recognition task [34]. Rats near-optimally maximized instantaneous reward rate (iRR) at the end of learning but chose response times that were too slow to be iRR-optimal early in learning. To understand the rats’ learning trajectory, we examined learning trajectories in a recurrent neural network (RNN) trained on the same task. We derive a reduction of this RNN to a learning drift diffusion model (LDDM) with time-varying parameters that describes the network’s average learning dynamics. Mathematical analysis of this model reveals a dilemma: at the beginning of learning when error rates are high, iRR is maximized by fast responses [10]. However, fast responses mean minimal stimulus exposure, little opportunity for perceptual processing, and consequently slow learning. Because of this learning speed/iRR (LS/iRR) trade-off, slow responses early in learning can yield greater total reward over engagement with the task, suggesting a normative basis for the rats’ behavior. We then experimentally tested and confirmed several model predictions by evaluating whether response time and learning speed are causally related, and whether rats choose their response times so as to take advantage of learning opportunities. Our results suggest that rats exhibit cognitive control of the learning process, adapting their behavior to approximately accrue maximal total reward across the entire learning trajectory, and indicate that a policy that prioritizes learning in perceptual tasks may be advantageous from a total reward perspective.

## Results

### Trained Rats Solve the Speed-Accuracy trade-off

We trained *n* = 26 rats on a visual object recognition two-alternative forced choice task (*see* Methods) [34]. The rats began a trial by licking the central of three capacitive lick ports, at which time a static visual object that varied in size and rotation from one of two categories appeared on a screen. After evaluating the stimulus, the rats licked the right or left lick port. When correct, they received a water reward, and when incorrect, a timeout period (Fig. 1a, Fig. S1). Because this was a free-response task, rats were also able to initiate a trial and not make a response, but these ignored trials made up a small fraction of all trials and were not considered during our analysis (Fig. S2).

**Figure 1:**
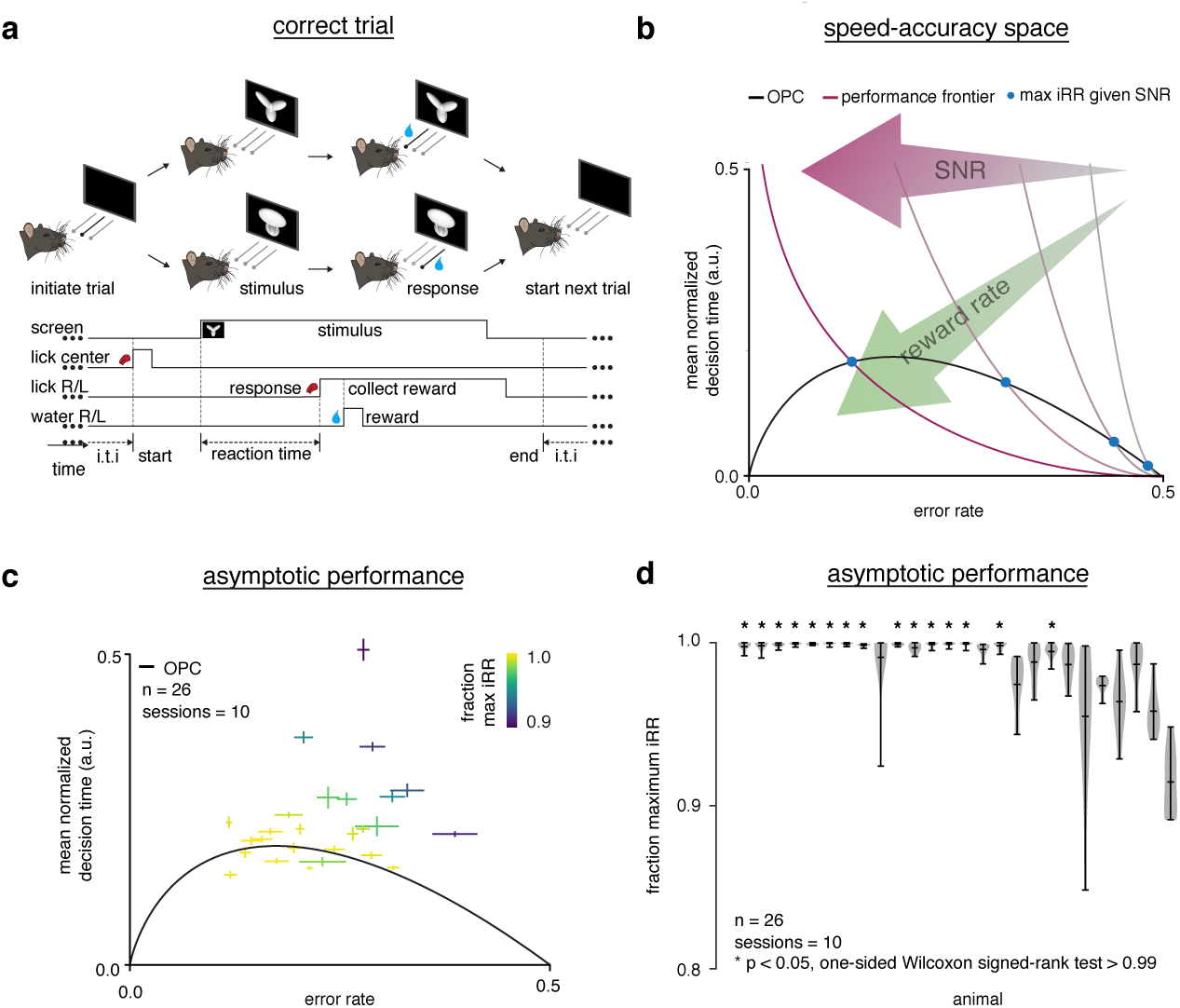
Trained rats solve the speed-accuracy trade-off. **(a)** Rat initiates trial by licking center port, one of two visual stimuli appears on the screen, rat chooses correct left/right response port for that stimulus and receives a water reward. **(b)** Speed-accuracy space: a decision making agent’s ER and mean normalized DT (a normalization of DT based on the average timing between one trial and the next, *see* Methods). Assuming a simple drift-diffusion process, agents that maximize iRR (*see* Methods: Evaluation of Optimality) must lie on an optimal performance curve (OPC, black trace) [10]. Points on the OPC relate error rate to mean normalized decision time, where the normalization takes account of task timing parameters (*e.g.* average response-to-stimulus interval). For a given SNR, an agent’s performance must lie on a performance frontier swept out by the set of possible threshold-to-drift ratios and their corresponding error rates and mean normalized decision times. The intersection point between the performance frontier and the OPC is the error rate and mean normalized decision time combination that maximizes iRR for that SNR. Any other point along the performance frontier, whether above or below the OPC, will achieve a suboptimal iRR. Overall, iRR increases toward the bottom left with maximal instantaneous reward rate at error rate = 0.0 and mean normalized decision time = 0.0. **(c)** Mean performance across 10 sessions for trained rats (*n* = 26) at asymptotic performance plotted in speed-accuracy space. Each cross is a different rat. Color indicates fraction of maximum instantaneous reward rate (iRR) as determined by each rat’s performance frontier. Errors are bootstrapped SEMs. **(d)** Fraction of maximum iRR, a quantification of distance to the OPC, for same rats and same sessions as **c**. Fraction of maximum iRR is a comparison of an agent’s current iRR with its optimal iRR given its inferred SNR. Approximately 15 of 26 (∼60%) of rats attain greater than 99% fraction maximum iRRs for their individual inferred SNRs. * denotes *p* < 0.05, one-tailed Wilcoxon signed-rank test for mean > 0.99.

We examined the relationship between error rate (ER) and decision time (DT) during asymptotic performance using the drift-diffusion model (DDM) (Fig. S3). In the DDM, perceptual information is integrated through time until the level of evidence for one alternative reaches a threshold. The speed-accuracy trade-off is controlled by the subject’s choice of threshold, and is solved when a subject’s performance lies on an optimal performance curve (OPC; Fig. 1b) [10]. The OPC defines the mean normalized DT and ER combination for which an agent will collect maximal iRR (*see* Methods). At any given time, an agent will have some perceptual sensitivity (signal-to-noise ratio, SNR) which reflects how much information about the stimulus arrives per unit time. Given this SNR, an agent’s position in speed-accuracy space (the space relating ER and DT) is constrained to lie on a performance frontier traced out by different thresholds (Fig. 1b). Using a low threshold yields fast but error-prone responses, while using a high threshold yields slow but accurate responses. An agent only maximizes iRR when it chooses the ER and DT combination on its performance frontier that intersects the OPC. After learning the task to criterion, over half the subjects collected over 99% of their total possible reward, based on inferred SNRs assuming a DDM (Fig. 1c, d).

Across a population, a uniform stimulus difficulty will reveal different SNRs because the internal perceptual processing ability in every subject will be different. Thus, although we did not explicitly vary stimulus difficulty [11, 18, 29, 30], as a population, animals clustered along the OPC across a range of ERs (Fig. 1d), supporting the assertion that well-trained rats achieve a near maximal iRR in this perceptual task. We note that subjects did not span the entire range of possible ERs, and that the differences in optimal DTs dictated by the OPC for the ERs we did observe are not large. It remains unclear whether our subjects would be optimal over a wider range of task parameters. Notwithstanding, previous work with a similar task found that rats did increase DTs in response to increased penalty times, indicating a sensitivity to these parameters [26]. Thus, for our perceptual task and its parameters, trained rats approximately solve the speed-accuracy trade-off.

### Rats Do Not Maximize Instantaneous Reward Rate During Learning

Knowing that rats harvested reward near-optimally after learning, we next asked whether rats harvested instantaneous reward near-optimally during learning as well. If rats optimized iRR throughout learning, their trajectories in speed-accuracy space should always track the OPC.

During learning, a representative individual (*n* = 1) started with long RTs that decreased as accuracy increased across training time (Fig. 2a). Transforming this trajectory to speed-accuracy space revealed that throughout learning the individual did not follow the OPC (Fig. 2b). Early in learning, the individual started with a much higher DT than optimal, but as learning progressed it approached the OPC. The maximum iRR opportunity cost is the fraction of maximum possible iRR relinquished for a choice of threshold (and average DT) (*see* Methods). We found that this individual gave up over 20% of possible iRR at the beginning of learning but harvested reward near-optimally at asymptotic performance (Fig. 2c). These trends held when the learning trajectories of *n* = 26 individuals were averaged (Fig. 2d-f). These results show that rats do not greedily maximize iRR throughout learning and lead to the question: if rats maximize iRR at the end of learning, what principle governs their strategy at the beginning of learning?

**Figure 2:**
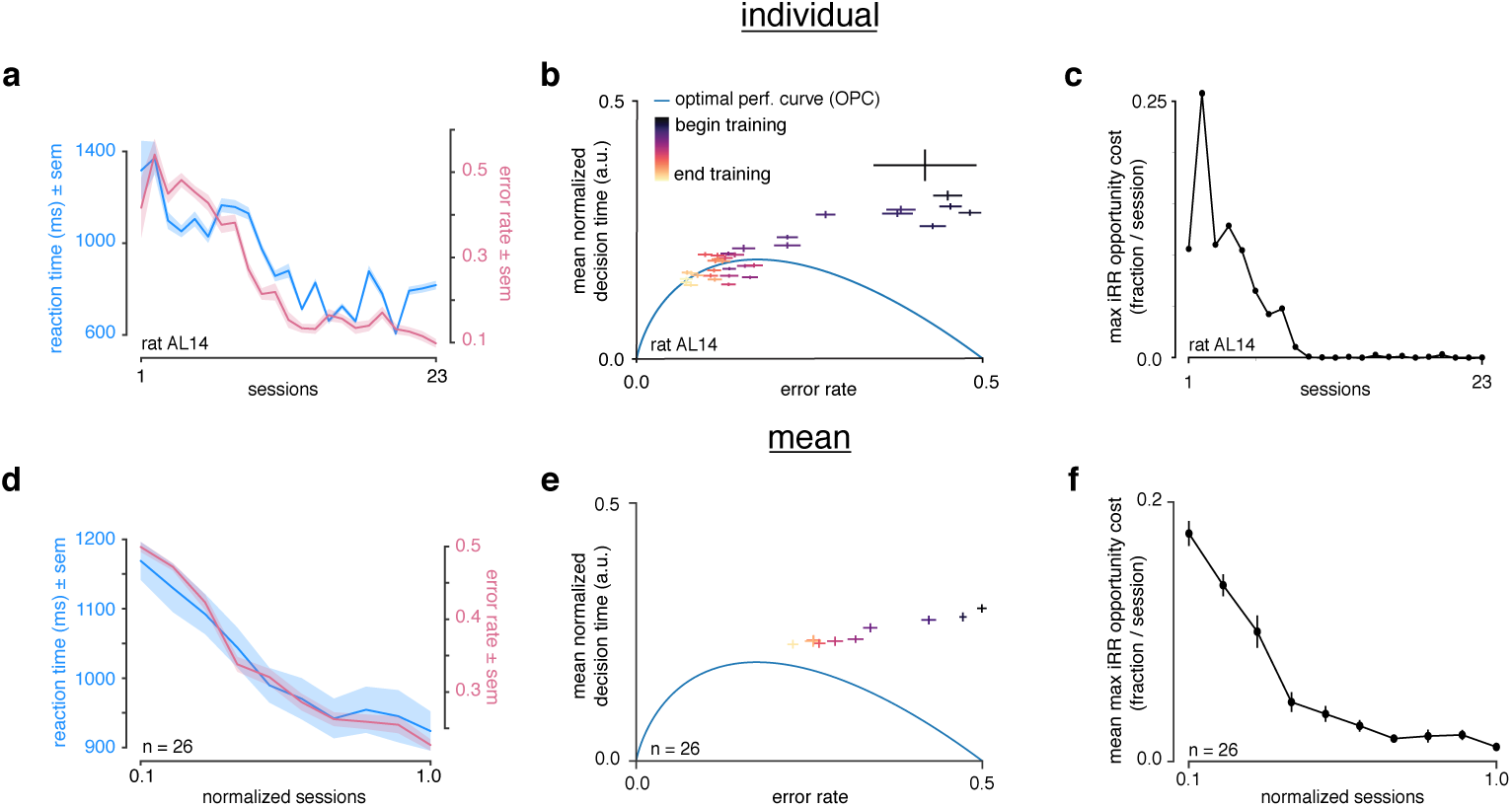
Rats do not greedily maximize instantaneous reward rate during learning. **(a)** Reaction time (blue) and error rate (pink) for an example subject (rat AL14) across 23 sessions. **(b)** Learning trajectory of individual subject (rat AL14) in speed-accuracy space. Color map indicates training time. OPC in blue. **(c)** Maximum iRR opportunity cost (*see* Methods) for individual subject (rat AL14) **(d)** Mean reaction time (blue) and error rate (pink) for *n* = 26 rats during learning. Sessions across subjects were transformed into normalized sessions, averaged and binned to show learning across 10 bins. Normalized training time allows averaging across subjects with different learning rates (*see* Methods). **(e)** Learning trajectory of *n* = 26 rats in speed-accuracy space. Color map and OPC as in **a. (f)** Maximum iRR opportunity cost of rats in **b** throughout learning. Errors reflect within-subject session SEMs for **a** and **b** and across-subject session SEMs for **d, e** and **f**.

### Model Reveals That Prioritizing Learning Can Maximize Total Reward

To simulate various learning strategies, we developed a simple linear recurrent neural network formalism for our task with the goal of investigating how long-term perceptual learning across many trials is influenced by the choice of DT (Box 1). A simple linear neural network processes input stimuli through weighted synaptic connections *w*. To model eye and body motion, the inputs are slightly jittered and rotated on each time step of a trial. For simplicity, the recurrent connectivity is fixed to 1 such that inputs are linearly integrated through time [35]. When the recurrent activity hits a threshold level, a response is made and the trial terminates. Then, to model perceptual learning, the perceptual weights *w* are updated using error-corrective learning, implemented as gradient descent on the hinge loss (*see* Methods).

While this model can be simulated to obtain sample learning trajectories, its average dynamics can also be solved analytically, yielding important insights. We derived a reduction of this model to a DDM with time-dependent parameters. This “learning DDM” (LDDM) closely tracked simulated trajectories of the full network (Box 1; Fig. S4; *see* Methods). In short, the reduction explains how SNR in a DDM model changes over time on average under error-corrective learning. In designing this model, we kept components as simple as possible to highlight key qualitative trade-offs between learning speed and decision strategy. Because of its simplicity, like the standard DDM, it is not meant to quantitatively describe all aspects of behavior. We instead use it to investigate qualitative features of decision making strategy, and expect that these features would be preserved in other related models of perceptual decision making [9, 36–48].

A key prediction of the LDDM is a tension between learning speed and iRR, the LS/iRR trade-off. This tension is clearest early in learning when ERs are near 50%. Then the rate of change in SNR is

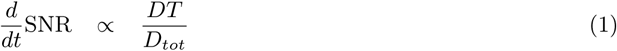

where the proportionality constant does not depend on *DT* (see derivation, Methods). Hence learning speed increases with increasing DT. By contrast the iRR when accuracy is 50% decreases with increasing DT. When encountering a new task, therefore, agents face a dilemma: they can either harvest a large iRR, or they can learn quickly.

**Box 1:**
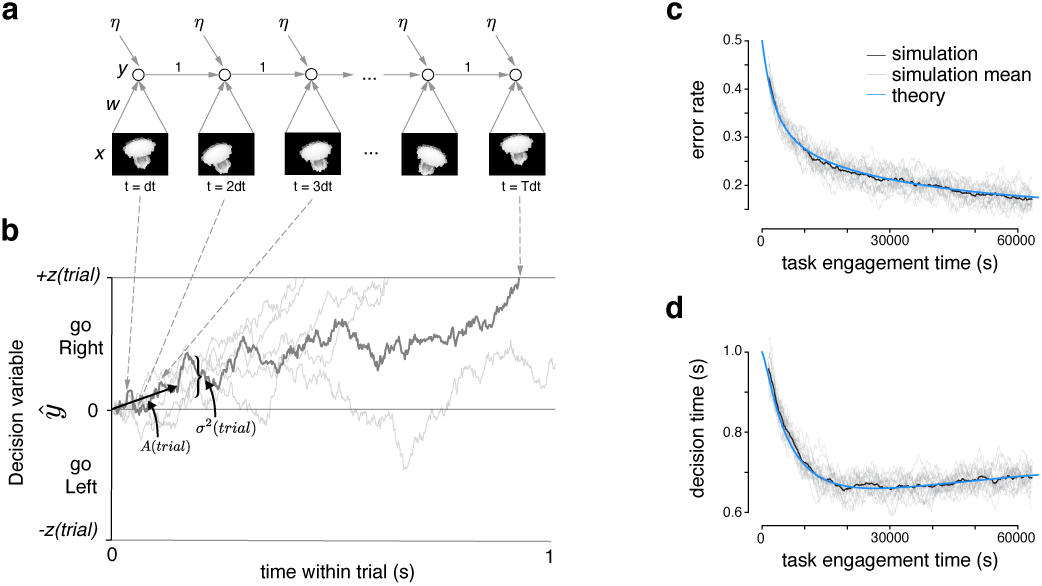
Recurrent Neural Network and Learning DDM models

Within a trial, inputs *x*(*t*) are filtered through perceptual weights *w*(*trial*) at discrete times *t* = 1*dt*, 2*dt*, … where *dt* is a small time step parameter, and added to a decision variable *ŷ*(*t*) along with i.i.d. integrator noise 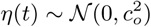. When the decision variable crosses a threshold ±*z*(*trial*), a decision is made. Panel A shows one example roll out of the recurrent neural network (RNN) through time. These within-trial dynamics in the limit of small time steps are equivalent to a standard drift-diffusion model (DDM) with a drift rate *A*(*trial*) toward the correct response and diffusion noise variance *σ*^2^(*trial*) that depend on the distribution of inputs *x*(*t*) and weights *w*(*trial*) (see derivation, Methods). Average performance is governed by the signal-to-noise ratio *Ā*(*trial*) = *A*(*trial*)^2^*/σ*^2^(*trial*) and threshold *z*(*trial*). Panel B shows the decision variable for the RNN (dark gray), and other trajectories of the equivalent DDM for different diffusion noise samples (light gray).

To implement perceptual learning, after each trial, weights in the recurrent network are updated using gradient descent on the hinge loss, corresponding to standard practice in deep learning:

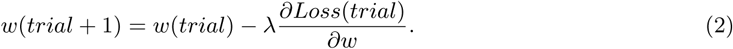

Here *λ* is a small learning rate and *Loss*(*trial*) = *max*(0, 1 − *y*(*trial*)*ŷ*(*trial*)) where *y*(*trial*) = ±1 is the correct output sign for the trial.

In the limit of small learning rates, applying Eq. (2) in the RNN is equivalent to the following SNR dynamics in the drift-diffusion model (see derivation, Methods):

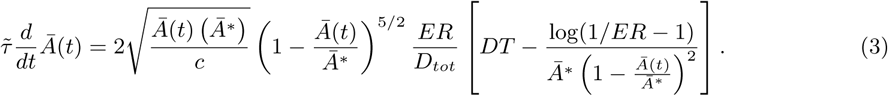

Time *t* measures seconds of task engagement. The SNR dynamics depend on five parameters: the time constant 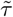 related to the learning rate, the initial SNR *Ā*(0), the asymptotic achievable SNR after learning *Ā*^*^, the integration-noise to input-noise variance ratio *c*, and the choice of threshold *z*(*t*) over training. Panels C-D show how *ER* and *DT* change over a long period of task engagement in the RNN (light gray, simulated individual traces; dark gray, mean) compared to the theoretical predictions from the learning DDM (blue).

An agent’s decision making strategy consists of their choice of threshold over time *z*(*t*). Threshold affects *DT* and *ER*, and through these, the learning dynamics in Eq. (3). We consider four threshold policies:

**iRR-Greedy** Threshold *z*^*g*^(*t*) is set to the threshold *z*^*^(*Ā*(*t*)) that maximizes instantaneous reward rate for current SNR, *z*^*g*^(*t*) = *z*^*^(*Ā*(*t*)).

**iRR-Sensitive** Threshold *z*^*s*^(*t*) decays with time constant *γ* from an initial value *z*^*s*^(0) = *z*_0_ toward the iRR-optimal threshold, 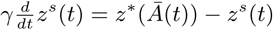.

**Constant** Threshold *z*^*c*^(*t*) is fixed to a constant *z*^*c*^(*t*) = *z*_0_.

**Global Optimal** Threshold *z*^*o*^(*t*) maximizes total reward over the duration of task engagement *T*_*tot*_,

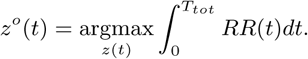

We approximately compute this threshold function using automatic differentiation (see Methods).

Just as the standard DDM instantiates different decision making strategies as different choices of threshold (for instance aimed at maximizing iRR, accuracy, or robustness) [30, 31], the LDDM instantiates different learning strategies through the choice of threshold trajectory over learning. In order to understand the rats’ learning trajectory, we evaluated several potential threshold policies (*see* Box 1 and Methods: Threshold Policies). An *iRR-greedy* threshold policy adjusts the threshold to maximize iRR at all times. This model is similar to a previously proposed neural network model of rapid threshold adjustment based on reward rate [16], and is iRR-optimal. A *constant* threshold policy is one where a target threshold that is optimal for some predicted future SNR is picked at the outset and kept constant throughout learning. Constant thresholds across difficulties have been found to be used as part of near-optimal and presumably cognitively cheaper strategies in humans [18]. In the *iRR-sensitive* threshold policy, the threshold starts at a specified initial value and then exponentially decays towards the threshold that would maximize iRR given the current SNR. Notably, with this policy, as the SNR changes due to learning, the target threshold also changes through time. Finally, in the *global optimal* threshold policy, the threshold trajectory throughout learning is optimized to maximize total cumulative reward at some known predetermined end to the task. We computed this globally optimal trajectory using automatic differentiation through a discretization of the reduction dynamics (*see* Methods). Because it relies on knowledge unavailable to the agent at the start of learning (such as the asymptotic achievable SNR after learning, and the total future duration of task engagement), this policy is not achievable in practice and serves as an idealized benchmark to which other models and the rats’ trajectory can be compared.

In order to qualitatively understand how these models behave through time, we visualized their learning dynamics. To approximately place the LDDM task parameters in a similar space to the rats, we performed maximum likelihood fitting using automatic differentiation through the discretized reduction dynamics (*see* Methods). The four policies we considered clustered into two groups, distinguished by their behavior early in learning. A “greedy” group, which contained just the iRR-greedy policy, remained always on the OPC (Fig. 3a), and had fast initial response times (Fig. 3b), a long initial period at high error (Fig. 3c), and high initial iRR (Fig. 3d). By contrast, a “non-greedy” group, which contained the iRR-sensitive, constant, and global optimal policies, started far above the OPC (Fig. 3a), and had slow initial response times (Fig. 3b), rapid improvements in ER (Fig. 3c), and low iRR (Fig. 3d). Notably, while members of the non-greedy group started off with lower iRR, they rapidly surpassed the slow learning group (Fig. 3d) and ultimately accrued more total reward (Fig. 3e). Overall, these results show that threshold strategy strongly impacts learning dynamics due to the learning speed/iRR trade-off (Fig. 3f), and that prioritizing learning speed can achieve higher cumulative reward than prioritizing instantaneous reward rate.

**Figure 3:**
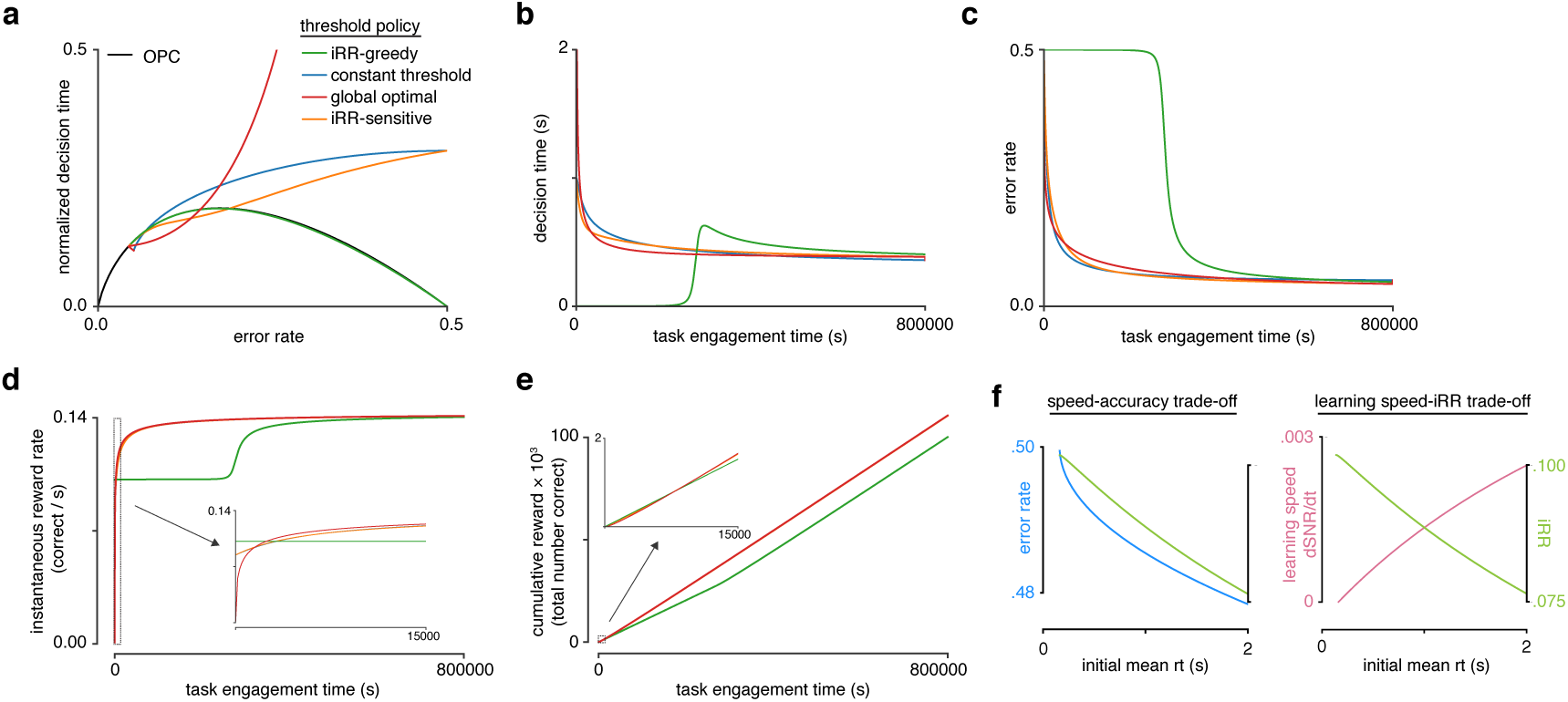
Model reveals rat learning dynamics lead to higher instantaneous reward rate and long-term rewards than greedily maximizing instantaneous reward rate. **(a)** Model learning trajectories in speed-accuracy space plotted against the OPC (black). **(b)** Decision time through learning for the four different threshold policies in **a. (c)** Error rate throughout learning for the four different threshold policies in **a. (d)** Instantaneous reward rate as a function of task engagement time for the full learning trajectory and a zoom-in on the beginning of learning (*inset*). **(e)** Cumulative reward as a function of task engagement time for the full learning trajectory and a zoom-in on the beginning of learning (*inset*). Threshold policies: iRR-greedy (green), constant threshold (blue), iRR-sensitive (orange) and global optimal (red). **(f)** In the speed-accuracy trade-off (*left*), ER (blue) decreases with increasing initial mean RT. iRR (green) at high error rates (∼0.5) also decreases with increasing initial mean RT. Thus, at high ERs, an agent solves the speed-accuracy trade-off by choosing fast RTs that result in higher ERs and maximize iRR. In the learning speed/iRR trade-off (*right*), initial learning speed (dSNR/dt, pink) increases with increasing initial mean RT, whereas iRR (green) follows the opposite trend. Thus, an agent must trade iRR in order to access higher learning speeds. Plots generated using LDDM model.

We further analyzed the differences between the three strategies in the non-greedy group. The global optimal policy selects extremely slow initial DTs to maximize the initial speed of learning. By contrast, the iRR-sensitive and constant threshold policies start with moderately slow responses. Nevertheless, we found that these simple strategies accrued 99% of the total reward of the global optimal strategy (Fig. S6). Hence these more moderate policies, which do not require oracle knowledge of future task parameters, derive most of the benefit in terms of total reward and may reflect a reasonable approach when the duration of task engagement is unknown.

Considering the rats’ trajectories in light of these strategies, their slow responses early in learning stand in stark contrast to the fast responses of the iRR-greedy policy (c.f. Fig. 2b, Fig. 3a). Equally, their responses were faster than the extremely slow initial DTs of the global optimal model. Both the iRR-sensitive and constant threshold models qualitatively matched the rats’ learning trajectory. However, DDM parameter fits of the rats’ behavior indicated that their thresholds decreased throughout learning, ruling out the constant threshold model (Fig. S5, *see* Methods). Furthermore, subsequent experiments (Fig. 4) also rule out a simple constant threshold strategy. Consistent with substantial improvements in perceptual sensitivity through learning, DDM fits to the rats also showed an increase in drift rate throughout learning (Fig. S5). Similar increases in drift rate have been observed as a universal feature of learning throughout numerous studies fitting learning data with the DDM [18, 49–53]. These qualitative comparisons suggest that rats adopt a “non-greedy” strategy that trades initial rewards to prioritize learning in order to harvest a higher iRR sooner and accrue more total reward over the course of learning.

**Figure 4:**
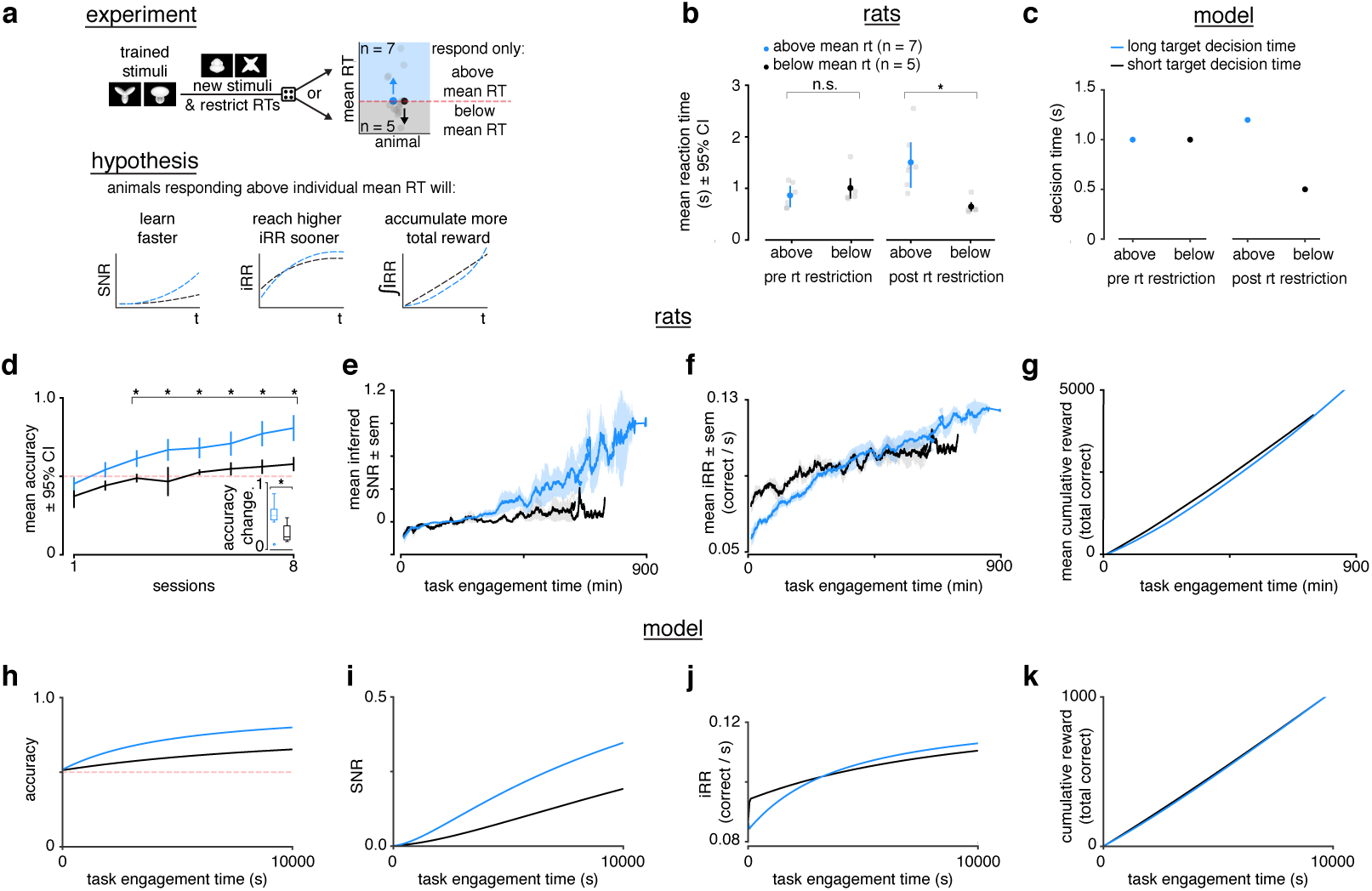
Longer reaction times lead to faster learning and higher instantaneous reward rates. **(a)** Schematic of experiment and hypothesized results. Previously trained animals were randomly divided into two groups: could only respond above (blue, *n* = 7) or below (black, *n* = 5) their individual mean reaction times for the previously trained stimulus and the new stimulus. Subjects responding above their individual mean reaction times were predicted to learn faster, reach a higher instantaneous reward rate sooner and accumulate more total reward. **(b)** Mean and individual reaction times before and after the reaction time restriction in rats. The mean reaction time for subjects randomly chosen to respond above their individual mean reaction times (blue, *n* = 7) was not significantly different to those randomly chosen to respond below their individual means (black, *n* = 5) before the restriction (Wilcoxon rank-sum test *p* > 0.05), but were significant after the restriction (Wilcoxon rank-sum test *p* < 0.05). Errors represent ±95% confidence intervals. **(c)** In the model a long (blue) and a short target decision time (black) were set. **(d)** Mean accuracy 95% confidence interval across sessions for rats required to respond above (blue, *n* = 7) or below (black, *n* = 5) their individual mean reaction times for a previously trained stimulus. Both groups had initial accuracy below chance because rats assume a response mapping based on an internal assessment of similarity of new stimuli to previously trained stimuli. To counteract this tendency and ensure learning, we chose the response mapping for new stimuli that contradicted the rats’ mapping assumption, having the effect of below-chance accuracy at first. * denotes *p* < 0.05 in two-sample independent t-test. *Inset* : accuracy change (slope of linear fit to accuracy across sessions to both groups, units: fraction per session). * denotes *p* < 0.05 in a Wilcoxon rank-sum test. **(e)** Mean inferred SNR, **(f)** mean iRR, and **(g)** mean cumulative reward across task engagement time for new stimulus pair for animals in each group. **(h)** Accuracy, **(i)** SNR, **(j)** iRR and **(k)** cumulative reward across task engagement time for long (blue) and short (black) target decision times in the LDDM.

### Learning Speed Scales with Reaction Time

To test the central prediction of the LDDM that learning (change in SNR) scales with mean DT, we designed a RT restriction experiment and studied the effects of the restriction on learning in the rats. Previously trained rats (*n* = 12) were randomly divided into two groups in which they would have to learn a new stimulus pair while responding above or below their individual mean RTs (‘slow’ and ‘fast’) for the previously trained stimulus pair (Fig. 4a). Before introducing the new stimuli, we carried out practice sessions with the new timing restrictions to reduce potential effects related to a lack of familiarity with the new regime. After the restriction, RTs were significantly different between the two groups (Fig. 4b). In the model, we simulated a RT restriction by setting two different DTs (Fig. 4c).

We found no difference in initial mean session accuracy between the two groups, followed by significantly higher accuracy in the slow group in subsequent sessions (Fig. 4d). The slope of accuracy across sessions was significantly higher in the slow group (Fig. 4d, *inset*). Importantly, the fast group had a positive slope and an accuracy above chance by the last session of the experiment, indicating this group learned (Fig. 4d).

Because of the speed-accuracy trade-off in the DDM, however, accuracy could be higher in the slow group even with no difference in perceptual sensitivity (SNR) or learning speed simply because on average they view the stimulus for longer during a trial, reflecting a higher threshold. To see if underlying perceptual sensitivity increased faster in the slow group, we computed the rats’ inferred SNR throughout learning, which takes account of the relationship between RT and ER. The SNR of the slow group increased faster (Fig. 4e), consistent with a learning speed that scales with DT.

We found that the slow group had a lower initial iRR, but that this iRR exceeded that of the fast group halfway through the experiment (Fig. 4f). Similarly, the slow group trended towards a higher cumulative reward by the end of the experiment (Fig. 4g). The LDDM qualitatively replicates all of our behavioral findings (Fig. 4h-k). These results demonstrate the potential total reward benefit of faster learning, which in this case was a product of enforced slower RTs.

Our experiments and simulations demonstrate that longer RTs lead to faster learning and higher reward for our task setting both *in vivo* and *in silico*. Moreover, they are consistent with the hypothesis that rats choose high initial RTs in order to prioritize learning and achieve higher iRRs and cumulative rewards during the task.

### Rats Choose Reaction Time Based on Learning Prospects

The previous experiments suggest that rats trade initial rewards for faster learning. Nonetheless, it is unclear how much control rats exert over their RTs. A control-free heuristic approach, such as adopting a fixed high threshold (our constant threshold policy), might incidentally appear near optimal for our particular task parameters, but might not be responsive to changed task conditions. If an agent is controlling the reward investment it makes in the service of learning, then it should only make that investment if it is possible to learn.

To test whether the rats’ RT modulations were sensitive to learning potential, we conducted a new experiment in which we divided rats into a group that encountered new learnable visible stimuli (*n* = 16, sessions = 13), and another that encountered unlearnable transparent or near-transparent stimuli (*n* = 8, sessions = 11) (Fig. 5a). From the perspective of the LDDM, both groups start with approximately zero SNR, however only the group with the visible stimuli can improve that SNR. If the rats choose their RTs based on how much it is possible to learn, then: (1) rats encountering stimuli that they can learn will increase their RTs to learn quickly and increase future iRR. (2) Rats encountering stimuli that they cannot learn might first increase their RTs to learn that there is nothing to learn, but (3) will subsequently decrease RTs to maximize iRR.

**Figure 5:**
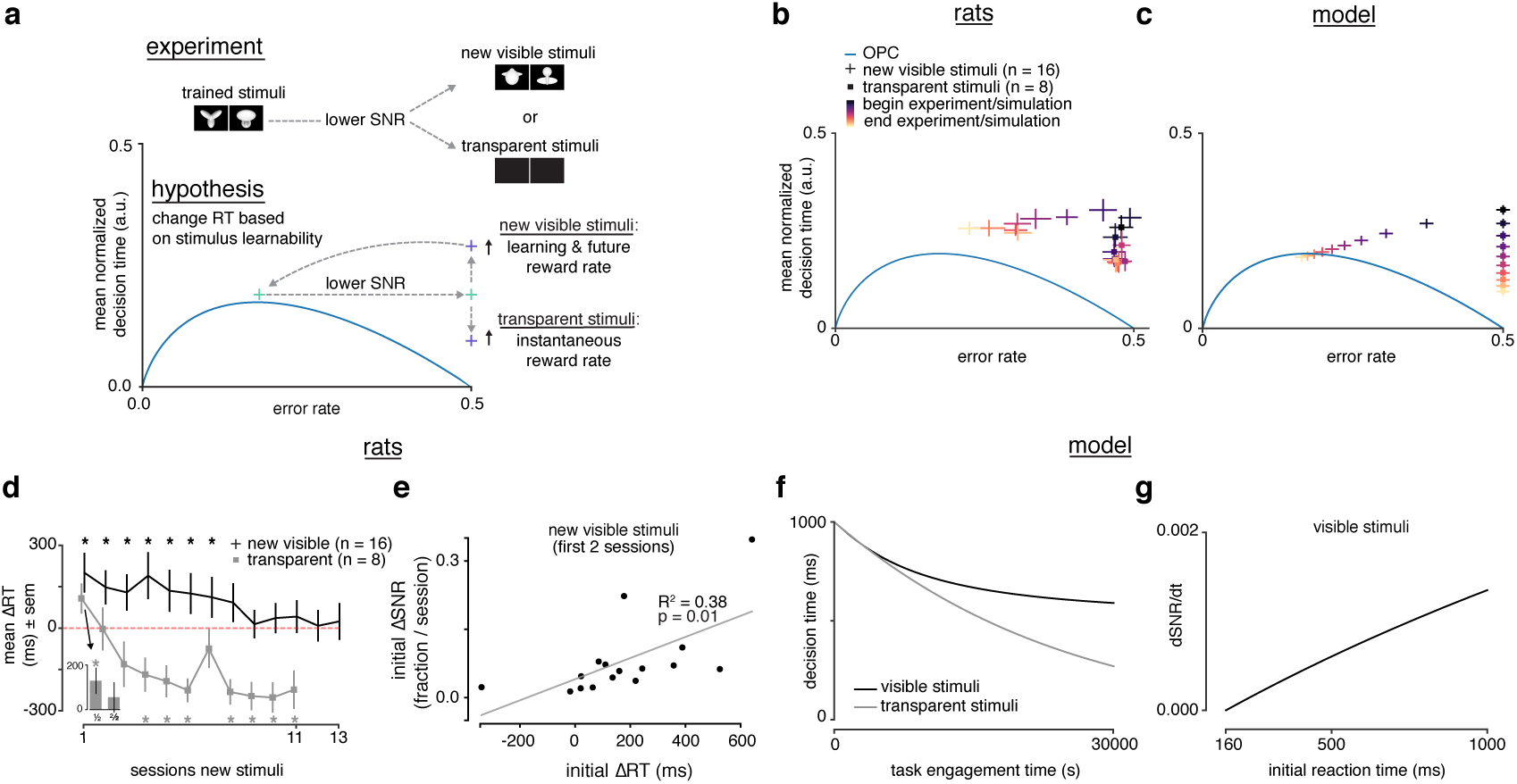
Rats choose reaction time based on stimulus learnability. **(a)** Schematic of experiment: rats trained on stimulus pair 1 were presented with new visible stimulus pair 2 or transparent (alpha = 0, 0.1) stimuli. If rats change their reaction times based on stimulus learnability, they should increase their reaction times for the new visible stimuli to increase learning and future iRR and decrease their reaction time to increase iRR for the transparent stimuli. **(b)** Learning across normalized sessions in speed-accuracy space for new visible stimuli (*n* = 16, crosses) and transparent stimuli (*n* = 8, squares). Color map indicates time relative to start and end of the experiment. **(c)** iRR-sensitive threshold model runs with “visible” (crosses) and “transparent” (squares) stimuli (modeled as containing some signal, and no signal) plotted in speed-accuracy space. The crosses are illustrative and do not reflect any uncertainty. Color map indicates time relative to start and end of simulation. **(d)** Mean change in reaction time across sessions for visible stimuli or transparent stimuli compared to previously known stimuli. Positive change means an increase relative to previous average. *Inset* : first and second half of first session for transparent stimuli. * denotes *p* < 0.05 in permutation test. **(e)** Correlation between initial individual mean change in reaction time (quantity in **d**) and change in SNR (learning speed: slope of linear fit to SNR per session) for first two sessions with new visible stimuli. R^2^ and *p* from linear regression in **d**. Error bars reflect standard error of the mean in **b** and **d. (f)** Decision time across time engagement time for visible and transparent stimuli runs in model simulation. **(g)** Instantaneous change in SNR (dSNR/dt) as a function of initial reaction time (decision time + non-decision time *t*_0_) in model simulation.

We found that the rats with the visible stimuli qualitatively replicated the same trajectory in speed-accuracy space that we found when rats were trained for the first time (Fig. 2b, Fig. 5b). Because these previously trained rats had already mastered the task mechanics, this result rules out non-stimulus-related learning effects as the primary explanation for long RTs at the beginning of learning. Any slowdown in RT in this experiment was only attributable to stimulus changes. We calculated the mean change in RT (mean ΔRT) of new stimuli versus known stimuli. The visible stimuli group had a significant slow-down in RT lasting many sessions that returned to baseline by the end of the experiment (Fig. 5d, black trace).

Rats with the transparent stimuli also approached the OPC by decreasing their RTs across sessions to better maximize iRR (Fig. 5b). After a brief initial slow-down in RT in the first half of the first session (Fig. 5d, *inset*), RTs rapidly decreased (Fig. 5d, grey trace). Notably, RTs fell below the baseline RTs, indicating a strategy of responding quickly, which is iRR-optimal for this zero SNR task. Hence rodents are capable of modulating their strategy depending on their learning prospects.

This experiment also argues against several simple strategies for choosing reaction times. If rats respond more slowly after error trials, a phenomenon known as post-error slowing (PES), they might exhibit slower RTs early in learning when errors are frequent [54]. Indeed, we found a slight mean PES effect of about 50ms that was on average constant throughout learning, though it was highly variable across individuals (Fig. S7). However, rats viewing transparent stimuli had ERs constrained to 50%, yet their RTs systematically decreased (Fig. 5b), such that PES alone cannot account for their strategy. Similarly, choosing RTs as a simple function of time since encountering a task would not explain the difference in RT trajectories between visible and transparent stimuli (Fig. 5d).

A simulation of this experiment with the iRR-sensitive threshold LDDM model qualitatively replicated the rats’ behavior (Fig. 5c, f, g). Rodent behavior is thus consistent with a threshold policy that starts with a relatively long DT upon encountering a new task, and then decays towards the iRR-optimal DT. All other threshold strategies we considered fail to account for the totality of the results. The iRR-greedy strategy—as before—stays pinned to the OPC and speeds up upon encountering the novel stimuli rather than slowing down. The constant threshold strategy fails to predict the speedup in DT for the transparent stimuli. Finally, the global optimal strategy (which has oracle knowledge of the prospects for learning in each task) behaves like the iRR-greedy policy from the start on the transparent stimuli as there is nothing to learn.

Our RT restriction experiment showed that higher initial RTs led to faster learning, a higher iRR and more cumulative reward. Consistent with these findings, there was a correlation between initial mean ΔRT and initial ΔSNR across subjects viewing the visible stimuli, indicating the more an animal slowed down, the faster it learned (Fig. 5e). We further tested these results in the voluntary setting by tracking iRR and cumulative reward for the rats in the learnable stimuli setting with the largest (blue, n = 4) and smallest (black, n = 4) “self-imposed” change in RT (Fig. 6a). The rats with the largest change started with a lower but ended with a higher mean iRR, and collected more cumulative reward (Fig. 6b, c). Thus, in the voluntary setting there is a clear relationship between RT, learning speed, and its total reward benefits.

**Figure 6:**
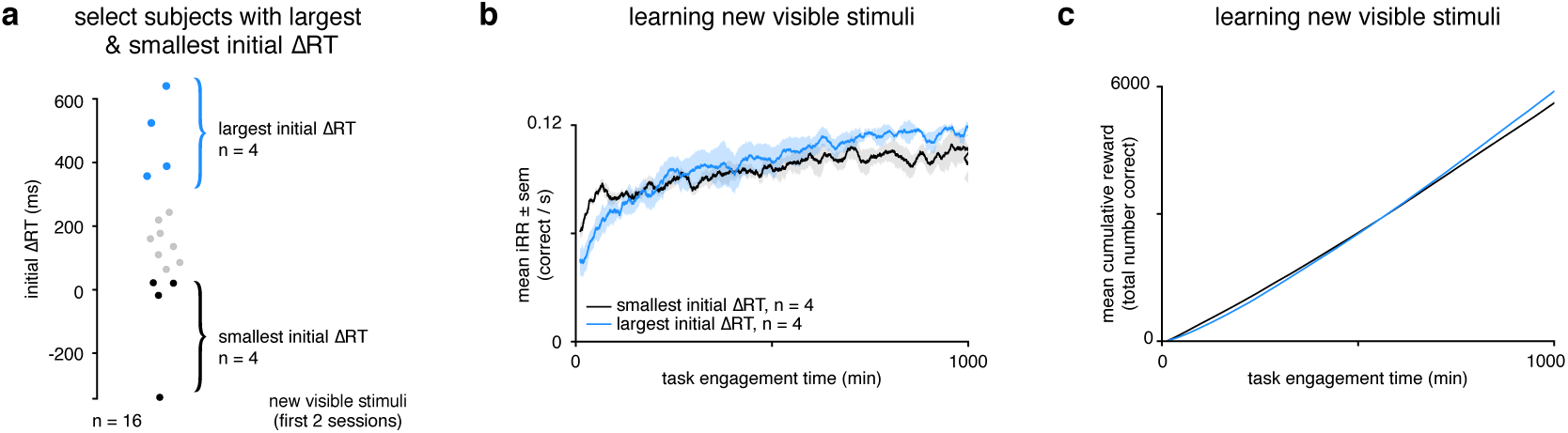
Rats that slowed down reaction times the most reached a higher instantaneous reward rate sooner and collected more reward. **a** Schematic showing segregation of top 25% of subjects (*n* = 4) with the largest initial ΔRTs for the new visible stimuli and the bottom 25% of subjects (*n* = 4) with the smallest initial ΔRTs. Initial ΔRTs were calculated as an average of the first 2 sessions for all subjects. **(b)** Mean iRR for subjects with largest and smallest mean changes in reaction time across task engagement time. **(c)** Mean cumulative reward over task engagement time for subjects as in **b**.

## Discussion

Our theoretical and empirical results identify a trade-off between the need to learn rapidly and the need to accrue immediate reward in a perceptual decision making task. We find that rats adapt their decision strategy to improve learning speed and approximately maximize total reward, effectively navigating this trade-off over the total period of task engagement. In our experiments, rats responded slowly upon encountering novel stimuli, but only when there was a visual stimulus to learn from, indicating they chose to respond more slowly in order to learn quickly. This behavior requires foregoing both a cognitively easier strategy—guessing as quickly as possible—and relinquishing a higher immediately available reward for several sessions spanning multiple days. By imposing different response times in groups of animals, we empirically verified our theoretical prediction that slow responses lead to faster learning and greater total reward in our task. These findings collectively show that rats exhibit cognitive control of the learning process, that is, the ability to engage in goal-directed behavior that would otherwise conflict with default or more immediately rewarding responses [22, 55–57].

Our high-throughput behavioral study with a controlled training protocol permits examination of the entire trajectory of learning, revealing hallmarks of non-greedy decision making. Nonetheless, it is accompanied by several experimental limitations. Our estimation of SNR improvements during learning relies on the drift-diffusion model. Importantly, while this approach has been widely used in prior work [18, 35, 49, 58], our conclusions are predicated on this model’s approximate validity for our task. Future work could address this issue by using a paradigm in which learners with different response deadlines are tested at the same fixed response deadline, equalizing the impact of stimulus exposure at test. This model-free paradigm is not trivial in rodents, because response deadlines cannot be rapidly instructed. Our study also focuses on one visual perceptual task. Further work should verify our findings with other perceptual tasks across difficulties, modalities, and organisms.

To understand possible learning trajectories, we introduced a theoretical framework based on a recurrent neural network, and from this derived a learning drift-diffusion model. The LDDM extends the canonical drift-diffusion framework to incorporate long-term perceptual learning, and formalizes a trade-off between learning speed and instantaneous reward. However, it remains approximate and limited in several ways. The LDDM builds off the simplest form of a drift-diffusion model, and various extensions and related models have been proposed to better fit behavioral data, including urgency signals [47, 59–62], history-dependent effects [63–69], imperfect sensory integration [35], confidence [58, 70, 71], and multi-alternative choices [72, 73]. More broadly, it remains unclear whether the drift-diffusion framework in fact underlies perceptual decision making, with a variety of other proposals providing differing accounts [38, 74, 75]. We speculate that the qualitative learning speed/instantaneous reward rate trade-off that we formally derive in the LDDM would also arise in other models of within-trial decision making dynamics. In addition, on a long timescale over many trials, the LDDM improves performance through error-corrective learning. Future work could investigate learning dynamics under other proposed learning algorithms such as feedback alignment [76], node perturbation [77], or reinforcement learning [33].

Conceptually, the learning speed/instantaneous reward rate trade-off is related to the explore/exploit trade-off common in reinforcement learning, but differs in detail. As traditionally framed in reinforcement learning, exploration permits sampling of the value associated with fully observable states and actions [78, 79]. By contrast, in our setting, the value of each action given object identity is clear, but object identity must be inferred from noisy measurements over time [80]. Exploration of the stimulus through long decision times permits integration of enough stimulus information such that error feedback reliably identifies informative stimulus features, improving future state inferences. Our trade-off therefore reflects a perceptual rather than value-based learning process. However, because there are reward benefits to improving future state inferences, this perceptual learning process is also necessarily value-based. The distinction between perceptual and value-based decision making may be due to the fact that they have been studied with paradigms obviating one another [81]. Even though our task was one with known decision values, the focus on learning revealed the value-based components of a perceptual learning process. Advances in deep reinforcement learning, where the marriage of perceptual learning mechanisms from deep learning and value-based learning mechanisms in reinforcement learning are an exciting avenue to further understand biological learning [82]. State-of-the-art deep reinforcement learning agents, which succeed in navigating the traditional explore/exploit dilemma on complicated tasks like Atari games, nevertheless fail to learn perceptual decisions like those considered here [83]. Our findings may offer routes to improving these artificial systems.

In order to navigate the learning speed/instantaneous reward rate trade-off, our findings suggest that rats deploy cognitive control of the learning process. Cognitive control has been defined as the allocation of mental faculties for goal-directed behavior over other default or more immediately rewarding behaviors [57]. Two main features of cognitive control govern its use: it is limited [84], and it is costly [55, 85–90]. If control is costly, then its application needs to be justified by the benefits of its application. The *Expected Value of Control (EVC)* theory posits that control is allocated in proportion to the expected value of control [56]. Previous work demonstrated that rats are capable of the economic reasoning required for optimal control allocation [91–93]. We demonstrated that rats incur a substantial initial instantaneous reward rate opportunity cost to learn the task more quickly, foregoing a cognitively less demanding fast guessing strategy that would yield higher initial rewards. Rather than optimizing instantaneous reward rate, which has been the focus of prior theories [9, 10, 18], our analysis suggests that rats approximately optimize total reward over task engagement. Relinquishing initial reward to learn faster, a cognitively costly strategy, is justified by a larger total reward over task engagement.

Assessing the expected value of learning in a new task requires knowing how much can be learned and how long the task will be performed. Neither of these quantities is directly observable upon first encountering a new task, opening the question of how rodents know to slow down in one task but not another. Importantly, rats only traded reward for information when learning was possible, a result in line with data demonstrating that humans are more likely to trade reward for information during long experimental time horizons, when learning is more likely [94]. Moreover, previous work has highlighted the explicit opportunity cost of longer deliberation times [47], a trade-off that will differ during learning and at asymptotic performance, as we demonstrate here. One possibility is that rats estimate learnability and task duration through meta-learning processes that learn to estimate the value of learning through experience with many tasks [95–97]. The amount of control allocated to learning the current task could be proportional to its estimated value, determined based on similarity to previous learning situations and their reward outcomes and control costs [98]. Previous observations of suboptimal decision times in humans analogous to those we observed in rats [11, 18, 30, 31], might reflect incomplete learning, or subjects who think they still have more to learn. Future work could test further predictions emerging from a control-based theory of learning. An agent should assess both the predicted duration of task engagement and the predicted difficulty of learning in order to determine the optimal decision making strategy early in learning, and this can be tested by, for instance, manipulating the time horizon and difficulty of the task.

The trend of a decrease in response time and an increase in accuracy through practice—which we observed in our rats—has been widely observed for decades in the skill acquisition literature, and is known as the *Law of Practice* [99–102]. Accounts of the Law of Practice have posited a cognitive control-mediated transition from shared/controlled to separate/automatic representations of skills with practice [101, 103–105]. On this view, control mechanisms are a limited, slow resource that impose unwanted processing delays. Our results suggest an alternative non-mutually-exclusive reward-based account for why we may so ubiquitously observe the Law of Practice. Slow responses early in learning may be the goal of cognitive control, as they allow for faster learning, and faster learning leads to higher total reward. When faced with the ever-changing tasks furnished by naturalistic environments, it is the speed of learning which may exert the strongest impact on total reward.

## Acknowledgments

We thank Chris Baldassano, Christopher Summerfield, Rahul Bhui and Grigori Guitchounts for useful discussions. We thank Joshua Breedon for summer assistance in developing faster animal training procedures. We thank Ed Soucy and the NeuroTechnology Core for help with improvements to the behavioral response rigs. This work was supported by the Richard A. And Susan F. Smith Family Foundation and IARPA contract # D16PC00002. J.M. was supported by the Harvard Brain Science Initiative (HBI) and the Department of Molecular and Cellular Biology at Harvard, and a Presidential Postdoctoral Research Fellowship at Princeton. A.M.S. was supported by a Swartz Postdoctoral Fellowship in Theoretical Neuroscience and a Sir Henry Dale Fellowship from the Wellcome Trust and Royal Society (Grant Number 216386/Z/19/Z).

## Author Contributions

J.M. and A.M.S. conceived the work, designed, ran and analyzed the experiments and simulations, and wrote the paper. D.D.C. provided input to experimental design. T.C. aided J.M. in establishing initial operant training procedures and behavioral analysis. J.Y.R. designed the behavioral response rigs.

## Methods

### Behavioral Training

#### Subjects

All care and experimental manipulation of animals were reviewed and approved by the Harvard Institutional Animal Care and Use Committee. We trained animals on a high-throughput visual object recognition task that has been previously described [34]. A total of 44 female Long-Evans rats were used for this study, with 38 included in analyses. Twenty-eight rats (AK1—12 & AL1—16) initiated training on stimulus pair 1, and 26 completed it (AK8 and AL12 failed to learn). Another 8 animals (AM1—8) were trained on stimulus pair 1 but were not included in the initial analysis focusing on asymptotic performance and learning (Fig. 1d, e; Fig. 2) because they were trained after the analyses had been completed. Subjects AM5—8, although trained, did not participate in other behavioral experiments so do not appear in this study. Sixteen animals (AL1—8, AL13—16 & AM1—8) participated in learning stimulus pair 2 (“new visible stimuli”; canonical only training regime) while 10 animals (AK1—3, 5—7, 9—12) initially participated in viewing transparent (alpha = 0; AK1, 3, 6, 7, 11) or near-transparent stimuli (alpha = 0.1; AK2, 5, 9, 10, 12), with the subjects sorted randomly into each group. The transparent and near-transparent groups were aggregated but 2 animals from the near-transparent group were excluded for performing above chance (AK5 & AK12) as this experiment focused on the effects of stimuli that could not be learned. The same 16 animals used for stimulus pair 2 were used for learning stimulus pair 3 under two different reaction time restrictions in which the subjects were sorted randomly. One rat (AL1) was excluded from the outset for not having learned stimulus pair 2. Two additional rats (AL4 & AL7) were excluded for not completing enough trials during practice sessions with the new reaction time restrictions. A final rat (AM1) was excluded because it failed to learn the task. The 12 remaining rats were grouped into 7 subjects required to respond above (AL3, AL8, AL13, AL15, AL16, AM3, AM4) and 5 subjects required to respond below their individual average reaction times (AL2, AL5, AL6, AL14, AM2). Finally, 8 rats (AN1—8) were trained on a simplified training regime (“canonical only”) used as a control for the typical “size & rotation” training object recognition regime (described below). Table S1 summarizes individual subject participation across behavioral experiments.

#### Behavioral Training Boxes

Rats were trained in high-throughput behavioral training rigs, each made up of 4 vertically stacked behavioral training boxes. In order to enter the behavioral training boxes, the animals were first individually transferred from their home cages to temporary plastic housing cages that would slip into the behavioral training boxes and snap into place. Each plastic cage had a porthole in front where the animals could stick out their head. In front of the animal in the behavior boxes were three easily accessible stainless steel lickports electrically coupled to capacitive sensors, and a computer monitor (Dell P190S, Round Rock, TX; Samsung 943-BT, Seoul, South Korea) at approximately 40° visual angle from the rats’ location. The three sensors were arranged in a straight horizontal line approximately a centimeter apart and at mouth-height for the rats. The two side ports (L/R) were connected to syringe pumps (New Era Pump Systems, Inc. NE-500, Farmingdale, NY) that would automatically dispense water upon a correct trial. The center port was connected to a syringe that was used to manually dispense water during the initial phases of training (see below). Each behavior box was equipped with a computer (Apple Macmini 6,1 running OsX 10.9.5 [13F34] or Macmini 7,1 running OSX El Capitan 10.11.13, Cupertino, CA) running MWorks, an open source software for running real-time behavioral experiments (MWorks 0.5.dev [d7c9069] or 0.6 [c186e7], The MWorks Project https://mworks.github.io/). The capacitive sensors (Phidget Touch Sensor P/N 1129 1, Calgary, Alberta, Canada) were controlled by a microcontroller (Phidget Interface Kit 8/8/8 P/N 1018 2) that was connected via USB to the computer. The syringe pumps were connected to the computer via an RS232 adapter (Startech RS-232/422/485 Serial over IP Ethernet Device Server, Lockbourne, OH). To allow the experimenter visual access to the rats’ behavior, each box was, in addition, illuminated with red LEDs, not visible to the rats.

#### Habituation

Long-Evans rats (Charles River Laboratories, Wilmington, MA) of about 250 g were allowed to acclimate to the laboratory environment upon arrival for about a week. After acclimation, they were habituated to humans for one or two days. The habituation procedure involved petting and transfer of the rats from their cage to the experimenter’s lap until the animals were comfortable with the handling. Once habituated to handling, the rats were introduced to the training environment. To allow the animals to get used to the training plastic cages, the feedback sounds generated by the behavior rigs, and to become comfortable in the behavior training room, they were transferred to the temporary plastic cages used in our high-throughput behavioral training rigs and kept in the training room for the duration of a training session undergone by a set of trained animals. This procedure was repeated after water deprivation, and during the training session undergone by the trained animals, the new animals were taught to poke their head out of a porthole available in each plastic cage to receive a water reward from a handheld syringe connected to a lickport identical to the ones in the behavior training boxes in the training rigs. Once the animals reliably stuck their head out of the porthole (one or two days) and accessed water from the syringe, they were moved into the behavior boxes.

#### Early Shaping

On their first day in the behavior boxes, rats were individually tutored as follows: Water reward was manually dispensed from the center lickport which is normally used to initiate a trial. When the animal licked the center lickport, a trial began. After a 500 ms tone period, one of two visual objects (stimulus pair 1) appeared on the screen (large front view, degree of visual angle 40°) chosen pseudo-randomly (three randomly consecutive presentations of one stimulus resulted in a subsequent presentation of the other stimulus). This appearance was followed by a 350 ms minimum reaction time that was instituted to promote visual processing of the stimuli. If the animal licked one of the side (L/R) lickports during this time, then the trial was aborted, there would be a minimum intertrial time (1300 ms), and the process would begin again.

At the time of stimulus presentation, a free water reward was dispensed from the correct side (L/R) lickport. If the animals licked the correct side lickport within the allotted amount of time (3500 ms) then an additional reward was automatically dispensed from that port. This portion of training was meant to begin teaching the animals the task mechanics, that is to first lick the center port, and then one of the two side ports.

After the rats were sufficiently engaged with the lickports and began self-initiating trials by licking the center lickport (usually 1 to several days, determined by experimenter) no more water was dispensed manually through the center lickport, but the free water rewards from the side lickports were still given. Once the rats were self-initiating enough trials without manual rewards from the center lickport (>200 per session), the free reward condition was stopped, and only correct responses were rewarded.

#### Training

Data collection for this study began once the rats had demonstrated proficiency of the task mechanics (as described above). The training curriculum followed was similar to that by Zoccolan and colleagues [34]. Rats performed the task for about 2 hours daily. Initially, the rats were only presented with large front views (40° visual angle, 0° of rotation) of the two stimuli (stimulus pair 1). Once the rats reached a performance level of ≥70% with these views, the stimuli decreased in size to 15° visual angle in a staircased fashion with steps of 2.5° visual angle. Once the rats reached 15° visual angle, rotations of the stimuli to the left or right were staircased in steps of 5° at a constant size of 30° visual angle. Once the rats reached ±60° of rotation, they were considered to have completed training and were presented with random transformations of the stimuli at different sizes (15° to 40° visual angle, step = 15°; 0° of rotation) or different rotations (−60 to +60° of rotation, step = 15°; 30° visual angle). After this point, ten additional training sessions were collected to allow the animals’ performance to stabilize with this expanded stimulus set.

During training, there was a bias correction that tracked the animals’ tendency to be biased to one side. If biased, stimuli mapped to the unbiased side were presented for a maximum of 3 consecutive trials. For example, if the bias correction detected an animal was biased to the right, the left-mapped stimulus would appear three trials at a time in a non-random fashion and the animals’ performance would drop from 50% to 25%, reducing the advantageousness of a biased strategy dramatically. If the animals continued to exhibit bias after one or two sessions of bias correction, then the limit was pushed to 5 consecutive trials. Once the bias disappeared, stimulus presentation resumed in a pseudo-random fashion.

The left/right mapping of the stimuli to lickports was counterbalanced across animals, ruling out any effects related left/right stimulus-independent biases, or left/right-independent stimulus bias across animals.

#### Training Regime Comparison

Although object recognition is supposed to be a fairly automatic process [106], it is possible that the 14 possible presentations of each stimulus of stimulus pair 1 (6 sizes at constant rotation, and 8 rotations at constant size) varied in difficulty. To rule out any possible difficulty effects during training and at asymptotic performance, We trained n = 8 different rats to asymptotic performance on the task but only on large, front-views of the visual objects (Fig. S8a). We compared the learning and asymptotic performance of the “size & rotation” cohort and the “canonical only” cohort across a wide range of behavioral measures. During learning, animals in both regimes followed similar learning trajectories in speed accuracy space (Fig. S8b), and clustered around the OPC at asymptotic performance (Fig. S8c). Comparisons of accuracy, reaction time, and fraction maximum instantaneous reward rate trajectories during learning and at averages asymptotic performance revealed no detectable differences (Fig. S8d—f). Total trials per session, and voluntary intertrial intervals after error trials did show slightly varied trajectories during learning, though there were no differences in their means after learning (Fig. S8g, h). The difference in total trials per session could be unrelated to the difference in training regimes. The difference in voluntary intertrial intervals, however, could be related to the introduction of different sizes and rotations: a sudden spike in this metric is seen about halfway through normalized sessions and decays over time. If this is the case, it is a curious result that rats choose to display their purported “surprise” in-between trials, and not during trials, as we found no difference in the reaction time trajectories. Both training regimes had overlapping fraction trials ignored metrics during learning, with a sharp decrease after the start, and a small significant difference in their number at asymptotic performance (Fig. S8i). We point out the fact that we do not consider voluntary intertrial intervals nor ignored trials in our analysis, so the differences between the regimes do not affect our conclusions. Overall, these results suggest that there is not a measurable or relevant difficulty effect based on our training regime with a variety of stimulus presentations.

#### Stimulus Learnability Experiment

##### Transparent Stimuli

In order to assess how animals behaved in a scenario with non-existent learning potential, a subset of already well-trained animals were presented with transparent (n = 5, alpha = 0) or near-transparent (n = 5, alpha = 0.1) versions of the familiar stimulus pair 1 for a duration of 11 sessions. Before these sessions, four sessions with stimulus pair 1 at full opacity (alpha = 1) were conducted to ensure animals could perform the task adequately before the manipulation. We predicted that the near-transparent condition would segregate animals into two groups, those that could perform the task and those that could not, based on each individual’s perceptual ability. The animals in the near-transparent condition that remained around chance performance (n = 3, rat AK2, AK9 & AK10) were grouped with the animals from the transparent condition, while those that performed well above chance (n = 2, rat AK5 & AK12) were excluded.

Reaction times were predicted to decrease during the course of the experiment, so to measure the change most effectively, the minimum reaction time requirement of 350 ms was removed. However, removing the requirement could lead to reduced reaction times regardless of the presented stimuli. To be able to measure whether the transparent stimuli led to a significant difference in reaction times compared to visible stimuli, we ran sessions with visible stimuli with no reaction time requirement for the same animals and compared these reaction times with those from the transparent condition. We found that the aggregate reaction time distributions were significantly different (Fig. S9a). A comparison of Vincentized reaction times revealed that there was a significant difference in the fastest reaction time decile (Fig. S9b), confirming that reaction times decreased significantly during presentation of transparent stimuli.

##### New Visible Stimuli

In order to assess how animals behaved in a scenario with high learning potential, a subset (n = 16) of already well-trained animals on stimulus pair 1 were presented with a never before seen stimulus pair (stimulus pair 2) for a duration of 13 sessions. Before these sessions, 5 sessions with the familiar stimulus pair 1 were recorded immediately preceding the stimulus pair 2 sessions in order to compare performance and reaction time after the manipulation for every animal. Previous pilot experiments showed that the animals immediately assigned a left/right mapping to the new stimuli based on presumed similarity to previously trained stimulus pair, so in order to enforce learning, the left/right mapping contrary to that predicted by the animals in the pilot tests was chosen. Because of this, animals typically began with an accuracy below 50%, as they first had to undergo reversal learning for their initial mapping assumptions. Because the goal of this experiment was to measure effects during learning and not demonstrate invariant object recognition, the new stimuli were presented in large front views only (visual angle = 40°, rotation = 0°).

### Behavioral Data Analysis

#### Software

Behavioral psychophysical data was recorded using the open-source MWorks 0.5.1 and 0.6 software (https://mworks.github.io/downloads/).The data were analyzed using Python 2.7 with the pymworks extension.

#### Drift-Diffusion Model Fit

In order to verify that our behavioral data could be modeled as a drift-diffusion process, the data were fit with a hierarchical drift-diffusion model [107], permitting subsequent analysis (such as comparison to the optimal performance curve) based on the assumption of a drift-diffusion process (Fig. S3). In order to assess parameter changes across learning, we fit the DDM to the start and end of learning for both stimulus pair 1 and stimulus pair 2, the first and second set of stimuli learned by the animals (Fig. S5). Drift rates increased and thresholds decreased by the end of learning, in agreement with previous findings [18, 49–53].

#### Behavioral Metrics

<u>Error Rate (ER)</u> was calculated by dividing the number of error trials by the number of total trials (error + correct) within a given window of trials or a the trials in a full behavioral training session. Accuracy was calculated as 1 - ER.

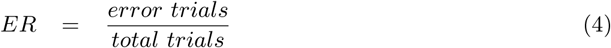

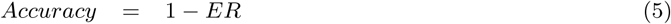

<u>Reaction Time (RT)</u> for one trial was measured by subtracting the time of the first lick on a response lickport from the stimulus onset time on the computer monitor. Mean RT was calculated by averaging reaction times across trials within a given window of trials or the trials in a full behavioral training session.

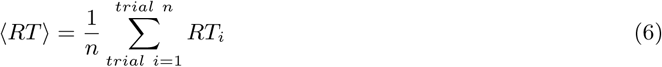

<u>Vincentized Reaction Time</u> is one method to report aggregate reaction time data meant to preserve individual distribution shape and be less sensitive to outliers in the group distribution [108, 109], although some scientists have argued parametric fitting (with an ex-Gaussian distribution, for example) and parameter averaging across subjects outperforms Vincentizing as sample size increases [110, 111]. Each subject’s reaction time distribution is divided into quantiles (*e.g.* deciles; similar to percentile, but between 0 and 1), and then the quantiles across subjects are averaged.

<u>Decision Time (DT)</u> for one trial was measured by subtracting the non-decision time T_0_ (*see* Estimating T_0_) from RT. Mean DT ⟨*DT*⟩ was calculated by subtracting T_0_ from the mean RT ⟨*RT*⟩across trials within a given window of trials or the trials in a full behavioral training session.

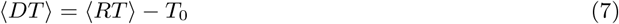

<u>Mean Normalized Decision Time ⟨</u>*<u>DT</u>*<u>⟩ /D_tot_</u> was measured by dividing mean DT ⟨*DT*⟩ by D_tot_, the sum of the non-decision time T_0_ and D_RSI_, the mean response-to-stimulus interval (*see* Determining D_RSI_, D_err_, D_corr_).

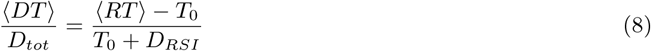

<u>Mean Difference in Mean Reaction Time ΔRT</u> was calculated by subtracting the mean reaction time of a number of baseline sessions from the mean reaction time of an experimental session. A positive difference indicates an increase over baseline mean reaction time. The mean of the two immediately preceding sessions with stimulus pair 1 were subtracted from the mean reaction time of every session with stimulus pair 2 or transparent stimuli for every animal individually (Fig. 5d, e). These differences were then averaged to get a mean difference in mean reaction time ΔRT.

<u>Mean Reward Rate ⟨</u>*<u>RR</u>*<u>⟩</u> is defined as mean accuracy per mean time per trial [9]:

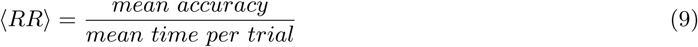

The mean time per trial is composed of the mean decision time ⟨*DT*⟩, non-decision-time T_0_, the post-error time D_err_ (*see* Determining D_RSI_, D_err_, D_corr_) scaled by the fraction of errors, and the post-correct time D_corr_ scaled by the fraction of correct choices:

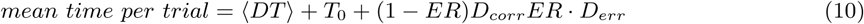

If we define D_p_ as the extra penalty time, that is, the difference between D_err_ and D_corr_:

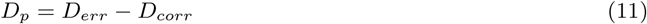

We can calculate the mean reward rate by equation A26 in Bogacz et al, 2006[10]:

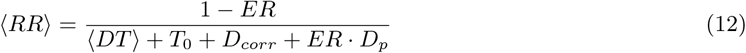

<u>Mean Total Correct Trials</u> is a model-free measure of the reward attained by the animals within a given window of trials. Every correct response yields an identical water reward, hence, reward can be counted by counting correct responses across trials. For one subject *a* ∈ [1, 2, 3,…, *k*], total correct trials at trial *n* are the sum of correct trials up to trial *n*:

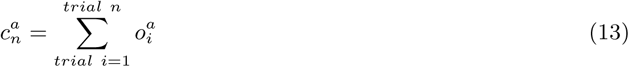

where 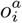 is an element in a vector *o*^*a*^ containing the outcomes of those trials 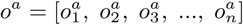. For correct and error responses 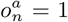 and 0 respectively (*e.g.* 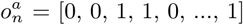).

Mean total correct trials up to trial *n* is calculated by taking the average of total correct trials across all animals *k* up to trial *n*.

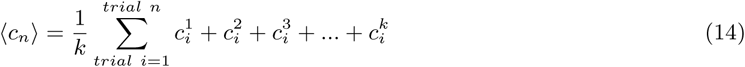

<u>Mean Cumulative Reward</u> is a measure of the reward attained by the animals within a given window of trials. To calculate this quantity, a moving average of RT and accuracy for a given window size are first calculated for every animal individually. To avoid averaging artifacts, only values a full window length from the beginning are considered. Given these moving averages, RR is then calculated for every animal and subsequently averaged across animals to get a moving average of mean reward rate. To calculate the mean cumulative reward, a numerical integral over a particular task time, such as *task engagement time* (*see* Measuring Task Time) is then calculated using the composite trapezoidal rule.

<u>Signal-to-Noise Ratio (SNR)</u> is a measure of an agent’s perceptual ability in a discrimination task. Given an animal’s particular ER and ⟨*DT*⟩, we use a standard drift-diffusion model equation to infer its SNR *Ā*:

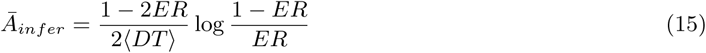

The SNR equation defines a U-shaped curve that increases as ERs move away from 0.5. For cases early in learning where ERs were below 0.5 because of potential initial biases, we assumed the inferred SNR was negative (meaning the animals had to unlearn the biases in order to learn, and thus had a monotonically increasing SNR during learning).

<u>SNR Performance Frontier</u> is a measure of an agent’s possible error rate and reaction time combinations based on their current perceptual ability. Because of the speed-accuracy trade-off, not all combinations of ER and ⟨*DT*⟩ are possible. Instead, performance is bounded by an agent’s SNR *Ā* at any point in time, and their particular (ER, ⟨*DT*⟩) combination will depend on their choice of threshold.

Given a fixed D_tot_ (as in the case of our experiment), this bound exists in the form of a performance frontier—-the combination of all resultant ERs and mean normalized DTs possible given a fixed SNR *Ā* and all possible thresholds 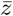.

Given an animal’s particular ER and ⟨*DT*⟩, we use a standard drift-diffusion model equation to infer its SNR *Ā*:

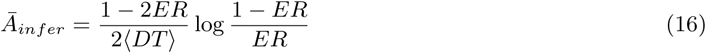

We can then use that *Ā*_*inf er*_ to calculate its performance frontier for a range of thresholds 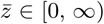 with standard equations from the drift-diffusion model:

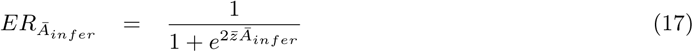

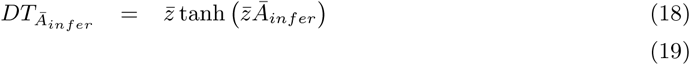

For every performance frontier there will be one unique 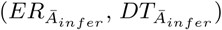 combination for which reward rate will be greatest, and it will lie on the OPC.

<u>Fraction Maximum Instantaneous Reward Rate</u> is a measure of distance to the optimal performance curve, *i.e.* optimal performance. Given an animal’s ER and ⟨*DT*⟩, we inferred their SNR and calculated their performance frontier as described above. We then divided the animal’s reward rate by the maximum reward rate on their performance frontier, corresponding to the point on the OPC they could have attained given their inferred SNR *A*_*infer*_:

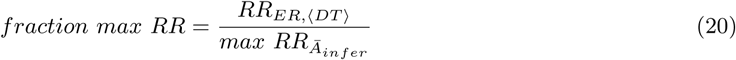

<u>Maximum Instantaneous Reward Rate Opportunity Cost</u>, like Fraction Maximum Reward Rate, is also measure of distance to the optimal performance curve, *i.e.* optimal performance, but it emphasizes the reward rate fraction given up by the subject given its current ER and ⟨*DT*⟩ combination along its SNR performance frontier. It is simply:

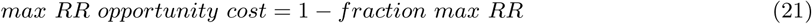

<u>Mean Post-Error Slowing</u> is a metric to account for the potential policy of learning by slowing down after error trials. In order to quantify the amount of post-error slowing in a particular subject, the subject’s reaction times for in a session are segregated into correct trials following an error, and correct trials following a correct choice, and separately averaged. The difference between these indicates the degree of post-error slowing present in that subject during that session.

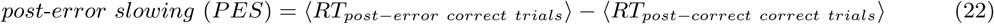

The mean post-error slowing for one session is thus the mean of this quantity across all subjects *k*.

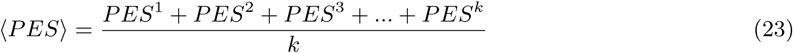

#### Computing Error

<u>Within-subject session errors</u> (*e.g.* Fig. 1d) for accuracy and reaction times were calculated by bootstrapping trial outcomes and reaction times for each session. We calculated a bootstrapped standard error of the mean by taking the standard deviation of the distribution of means from the bootstrapped samples. A 95% confidence interval can be calculated from the distribution of means as well.

<u>Across-subject session errors</u> (*e.g.* Fig. 5d) were computed by calculating the standard error of the mean of individual animal session means.

<u>Across-subject sliding window errors</u> (*e.g.* Fig. 5b; Fig. 6b) were calculated by averaging trials over a sliding window (*e.g.* 200 trials) for each animal first, then taking the standard error of the mean of each step across animals. Alternatively, the average could be taken across a quantile (*e.g.* first decile, second decile, etc.), and then the standard error of the mean of each quantile across animals was computed.

#### Measuring Task Time

<u>Trials</u> are the smallest unit of behavioral measure in the task and are defined by one stimulus presentation accompanied by one outcome (correct, error) and one reaction time.

<u>Sessions</u> are composed of as many trials as an animal chooses to complete within a set window of wall clock time, typically around 2 hours once daily. An error rate (fraction of error trials over total trials for the session) and a mean reaction time can be calculated for a session.

<u>Normalized Sessions</u> are a group of sessions (*e.g.* 1, 2, 3, …, 10) where a particular session’s normalized index corresponds to its index divided by the total number of sessions in the group (*e.g.* 0.1, 0.2, 0.3, …, 1.0). Because animals may take different numbers of sessions in order to learn to criterion, for instance, a normalized index for sessions allows better comparison of psychophysical measurements throughout learning.

<u>Stimulus Viewing Time</u> measures the time that the animals are viewing the stimulus, defined as the sum of all reaction times up to trial *n* as:

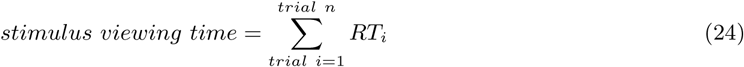

<u>Task Engagement Time</u> measures the time relevant for reward rate. We define task engagement time as the sum of reaction times plus all mandatory task time, essentially the cumulative sum of the average task time used to calculate reward rate:

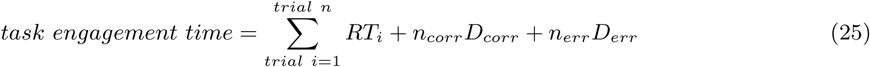

the sum of reaction times up to trial *n* plus the sum of D_err_ = 3136 ms and D_corr_ = 6370 ms, the mandatory post-error and post-correct response-to-stimulus intervals, proportional to the number of error and correct trials (*n = n*_*corr*_ + *n*_*err*_).

#### Statistical Analyses

<u>Figure 1d</u>: We wished to test whether the mean fraction maximum reward rate of our subjects over the ten sessions after having completed training were significantly different from optimal performance. A Shapiro-Wilk test failed to reject (p<0.05) a null hypothesis for normality for 18/26 subjects, with the following p-values (from left to right): (0.8162, 0.1580, 0.3746, 0.6985, 0.0025, 0.0467, 0.0040, 0.6522, 0.0109, 0.1625, 1.8178e-05, 0.0901, 0.7606, 0.0295, 0.0009, 0.2483, 0.5627, 0.0050, 0.4464, 0.6839, 0.5953, 0.0140, 0.1820, 0.1747, 0.6385, 0.2304). Thus, we conducted a one-sided Wilcoxon signed-rank test on our sample against 0.99, testing for the evidence that each subject’s mean fraction max reward rate was greater than 99% of the maximum (p<0.05), and obtained the following p-values (from left to right): (0.0025, 0.0025, 0.0025, 0.1013, 0.2223, 0.0063, 0.0047, 0.0025, 0.0025, 0.0025, 0.0025, 0.0571, 0.6768, 0.0047, 0.7125, 0.0372, 0.8794, 0.4797, 0.7125, 0.8987, 0.0372, 0.0109, 0.9975, 0.9766, 0.9917, 0.9975).

<u>Figure 4b</u>: We wished to test the difference in mean RT between two randomly chosen groups of animals before and after a RT restriction to assess the effectiveness of the restriction. A Shapiro-Wilk test did not support an assumption of normality for the ‘below’ group in either condition resulting in the following (W statistic, p-value) for the pre-RT restriction ‘above’ and ‘below’ groups and post-RT restriction ‘above’ and ‘below’ groups: (0.9073, 0.3777), (0.6806, 0.0059), (0.8976, 0.3168), (0.6583, 0.0033). Hence, we conducted a Wilcoxon rank-sum test for the pre- and post-RT restriction groups and found the pre-RT restriction group was not significant (p = 0.570) while the post-RT restriction group was (p = 0.007), indicating the two groups were not significantly different before the RT restriction, but became significantly different after the restriction.

<u>Figure 4d</u>: We wished to test the difference in accuracy between the ‘above’ and ‘below’ groups for every session of stimulus pair 3. A Shapiro-Wilk Test failed to reject the assumption of normality (p < 0.05) for any session from either condition (except session 4, ‘above’, which could be expected given there were 16 tests), with the following (W statistic, p-value) for [session: ‘above’, ‘below’] by session: [1: (0.9340, 0.6240),(0.8959, 0.3068)], [2: (0.9381, 0.6522), (0.8460, 0.1130)], [3: (0.9631, 0.8291), (0.9058, 0.3676)], [4: (0.7608, 0.0374), (0.9728, 0.9177)], [5: (0.8921, 0.3680), (0.9779, 0.9486)], [6: (0.7813, 0.0565), (0.9702, 0.9002)], [7: (0.8942, 0.3786), (0.9711, 0.9062)], [8: (0.7848, 0.0605), (0.9611, 0.8280)]

A Levene Test failed to reject the assumption of equal variances for every pair of sessions except the first (statistic, p-value): (6.3263, 0.0306), (2.2780, 0.1621), (1.2221, 0.2948), (0.8570, 0.3764), (2.7979, 0.1253), (0.7364, 0.4109), (0.0871, 0.7739), (0.0088, 0.9269).

Hence, we performed a two-sample independent *t* -test for every session with the following p-values: (0.4014, 0.04064, 0.0057, 0.0038, 0.0011, 0.0038, 0.0006, 6.3658e-05).

We also wished to test the difference between the slopes of linear fits to the accuracy curves for both conditions. A Shapiro-Wilk Test failed to reject the assumption of normality (p < 0.05) for either condition, with the following (W statistic, p-value) for ‘above’ and ‘below’: (0.8964, 0.3095), (0.8794, 0.3065). A Levene Test failed to reject the assumption of equal variances (p < 0.05) for each condition (statistic, p-value): (0.2141, 0.6535). Hence, we performed a two-sample independent *t* -test and found a significant difference (p = 0.0027).

<u>Figure 5d, e:</u> We wished to test whether the animals had significantly changed their session mean RTs with respect to their individual previous baseline RTs (paired samples). To do this, we conducted a permutation test for every session with the new visible stimuli (stimulus pair 2) or the transparent stimuli. For 1000 repetitions, we randomly assigned labels to the experimental or baseline RTs and then averaged the paired differences. The p-value for a particular session was the fraction of instances where the average permutation difference was more extreme than the actual experimental difference. For sessions with stimulus pair 2, the p-values from the permutation test were: (0.0034, 0.0069, 0.0165, 0.0071, 0.0291, 0.0347, 0.06, 0.0946, 0.3948, 0.244, 0.244, 0.4497, 0.3437). For sessions with transparent stimuli (plus rats AK2, AK9, & AK10 from the near-transparent stimuli) the p-values from the permutation were (0.0859375, 0.44921875, 0.15625, 0.03125, 0.02734375, 0.015625, 0.26953125, 0.02734375, 0.03125, 0.01953125, 0.0546875). To investigate whether the animals’ significantly slowed down their mean RTs compared to baseline during the first session of transparent stimuli, we divided RTs in the first session in half and ran a permutation test on each half with the following p-values: (0.0390625, 0.2890625).

In order to test the correlation between the initial change in RT and the initial change in SNR for stimulus pair 2, we ran a standard linear regression on the average per subject for each of these variables for the first 2 sessions of stimulus pair 2. The R^2^ refers to the square of the correlation coefficient, and the p-value is from a Wald Test with *t* -distribution of the test statistic.

<u>Figure S5a—h</u>: Statistical significance of differences in means between start of learning and after learning for stimulus pair 1 for the average predicted threshold distribution, and for the individual predicted thresholds were determined via a Wilcoxon signed-rank test. The p-values were: (a) average predicted threshold: <1e-4, (b) individuals predicted threshold: 0.0006, (c) average predicted drift rate: <1e-4, (d) individuals predicted drift rate: <1e-4.

For stimulus pair 2 we also used a Wilcoxon signed-rank test, and performed three tests per condition: baseline stimulus pair 1 versus start learning, start learning versus after learning, and baseline stimulus pair 1 versus after learning. In this order, the p-values were: (a) average predicted threshold: <1e-4, <1e-4, 0.3016, (b) individuals predicted threshold: 0.0386, 0.0557, 0.8767, (c) average predicted drift rate: <1e-4, <1e-4, <1e-4, (d) individuals predicted drift rate: 0.0004, 0.0004, 0.1089.

<u>Figure S7b, d:</u> We tested for a difference in mean post-error slowing between the first 2 sessions and last 2 sessions of training for each animal for stimulus pair 1 (b) or the last 2 sessions of stimulus pair 1 and the first 2 sessions of stimulus pair 2 (d) via a Wilcoxon-signed rank test. The p-values were (b) 0.585 and (d) 0.255.

<u>Figure S8d—i</u>: Statistical significance of differences in means between the two training regimes for a variety of psychophysical measures was determined by a Wilcoxon rank-sum test with p ¡ 0.05. The p-values were: (d) accuracy: 0.21, (e) reaction time: 0.81, (f) fraction max iRR: 0.22, (g) total trial number: 0.46, (h): voluntary iti after error: 0.75, (i) fraction trials ignored: 0.03.

<u>Figure S9a, b:</u> We tested for a difference in the aggregate reaction time distributions of a transparent stimuli condition (n = 5 subjects), and a no minimum reaction time condition with known stimuli (n = 5 subjects) via a 2-sample Kolmogorov-Smirnov Test and found a p-value of <1e-10.

We tested for the difference in the first decile of vincentized reaction times between these two conditions via a Wilcoxon rank-sum test and found a p-value of 0.047.

#### Evaluation of Optimality

Under the assumptions of a simple drift-diffusion process, the optimal performance curve (OPC) defines a set of optimal threshold-to-drift ratios with corresponding decision times and error rates for which an agent maximizes instantaneous reward rate [10]. Decision times are scaled by the particular task timing as mean normalized decision time: ⟨*DT*⟩ /D_tot_. The OPC is parameter free and can thus be used to compare performance across tasks, conditions, and individuals. An optimal agent will lie on different points on the OPC depending on differences in task timing (D_tot_) and stimulus difficulty (SNR). Assuming constant task timing, the SNR will determine different positions along the OPC for an optimal agent. For ⟨*DT*⟩ > 0 and 0 < ER < 0.5, the OPC is defined as:

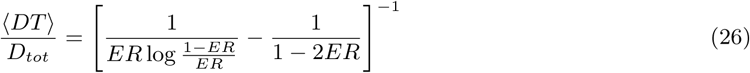

and exists in **speed-accuracy space**, defined by ⟨*DT*⟩ */D*_*tot*_ and ER. Given estimates for T_0_ and D_RSI_, the *ER* and ⟨*DT*⟩ for any given animal can be compared to the optimal values defined by the OPC in speed-accuracy space.

Moreover, because ER should decrease with learning, learning trajectories for different subjects and models can also be compared to the OPC and to each other in speed-accuracy space.

#### Mean Normalized Decision Time Depends Only on T_0_ and D_err_

To determine what times the normalizing term D_tot_ includes, we re-derive the OPC from average reward rate. According to Gold & Shadlen [9], average reward rate is defined as:

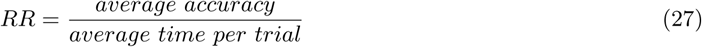

The average time per trial is composed of the average decision time, non-decision-time, the post-error time scaled by number of errors, and the post-correct time scaled by the number of correct choices:

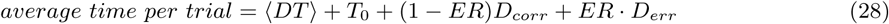

Define the extra penalty time *D*_*p*_ = *D*_*err*_ − *D*_*corr*_. We can write the average reward rate as equation A26 from [10],

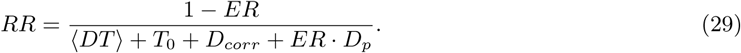

Optimal behavior is defined as maximizing reward rate with respect to the thresholds in the drift-diffusion model. We thus re-write ER and DT in terms of average threshold and average SNR,

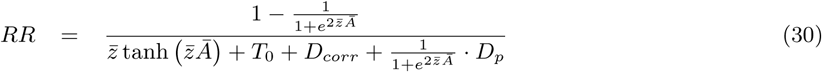

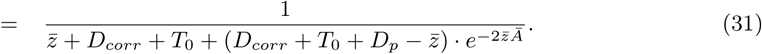

Next to find the extremum, we take the derivative of *RR* with respect to the threshold and set it to zero,

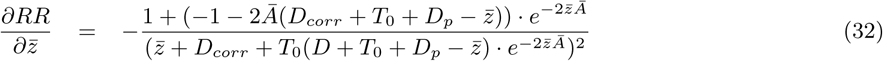

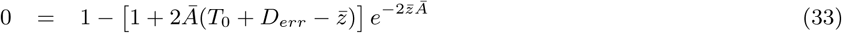

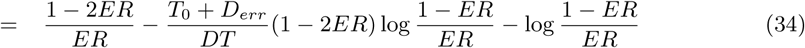

where in the final step we have rewritten 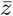 and *Ā* in terms of ER and ⟨*DT*⟩ .

Rearranging to place DT on the left hand side reveals an OPC where decision time is normalized by the post-error response-to-stimulus time *D*_*err*_ :

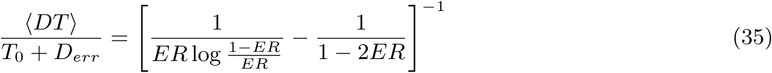

Notably, the post-correct response-to-stimulus time *D*_*corr*_ is not part of the normalization. Intuitively, this is because post-correct delays are an unavoidable part of accruing reward and therefore do not influence the optimal policy.

#### Estimating T_0_

*T*_*0*_ is defined as the non-decision time component of a reaction time, comprising motor and perceptual processing time [31]. It can be estimated by fitting a drift-diffusion model to the psychophysical data. Because of the experimentally imposed minimum reaction time meant to ensure visual processing of the stimuli, however, our reaction time distributions were truncated at 350 ms, meaning a drift-diffusion model fit estimate of T_0_ is likely to be an overestimate. To address this issue, we set out to determine possible boundaries for T_0_ and estimated it in a few ways, all of which did indeed fall between those boundaries (Fig. S10e).

We found that after training, in the interval between 350-375 ms, nearly all of our animals had accuracy measurements above chance (Fig. S10b), meaning that the minimum reaction time of 350 ms served as an upper bound to possible T_0_ values.

To determine a lower bound, we obtained measurements for the two components comprising T_0_: motor and initial perceptual processing times. To measure the minimum motor time required to complete a trial, we analyzed licking times across the different lickports. The latency from the last lick in the central port to the first lick in one of the two side ports peaked at around 80 ms (Fig. S10c). In addition, the latency from one lick to the next lick at the same port at any of the lickports was also around 80 ms (data not shown). Thus, the minimum motor time was determined by the limit on licking frequency, and not on a movement of the head redirecting the animal from the central port to one of the side ports. To measure the initial perceptual processing times, we looked to published latencies of visual stimuli traveling to higher visual areas in the rat. Published latencies reaching area TO (predicted to be after V1, LM and LI in the putative ventral stream in the rat) were around 80 ms (Fig. S10d) [112]. Based on these measurements, we estimated a T_0_ lower bound of approximately 160 ms.

One worry is that our lower bound could potentially be too low, as it is only estimated indirectly. Recent work on the speed-accuracy trade-off in a low-level visual discrimination tasks in rats found that accuracy was highest at a reaction time of 218 ms [28]. However, accuracy was still above chance for reaction times binned between 130—180 ms. In this task, reaction time was measured when an infrared beam was broken, which means we can assume there was no motor processing time. This leaves decision time, and initial perceptual processing time (part of T_0_) within the 130-180 ms duration. The complexity of solving a high-level visual task like ours and a low-level one will result in substantial differences in decision time, but should not in principle affect non-decision time. Considering a latency estimate of 80 ms based on physiological evidence [112] can account for the initial perceptual processing component of T_0_ and gives an estimate T_0_ = 80 ms for this study.

Because a reaction time around T_0_ should not allow for any decision time, accuracy should be around 50%. To estimate T_0_ based on this observation, we extrapolated the time at which accuracy would drop to 50% after plotting accuracy as a function of reaction time (Fig. S10a) and found values of 165 and 225 ms for linear and quadratic extrapolations respectively. Finally, we fit our behavioral data with a hierarchical drift-diffusion model [107] and found a T_0_ estimate of 295 ± 4 ms (despite there being no data below 350 ms). To address this issue, we fit drift-diffusion model to a small number of behavioral sessions we conducted with animals trained on the minimum reaction time of 350 ms but where that constraint was eliminated and found a T_0_ estimate of 265 ± 120 (SD) ms. We stress that because the animals were trained with a minimum reaction time, they likely would have required extensive training without that constraint to fully make use of the time below the minimum reaction time, thus this estimate is likely to also be an overestimate. We do note however that the estimate is lower than the estimate with an enforced minimum reaction time and has a much higher standard deviation (spanning our lower and upper bound estimates).

Despite the range of possible T_0_ values, we find that our qualitative findings (in terms of learning trajectory and near-optimality after learning) do not change (Fig. S10f, g), and proceed with a T_0_ = 160 ms for the main text.

#### Determining D_err_ and D_corr_

The experimental protocol defines the mandatory post-error and post-correct response-to-stimulus times (D_err_ and D_corr_ respectively). However, these times may not be accurate because of delays in the software communicating with different components such as the syringe pumps, and other delays such as screen refresh rates. We thus determined the actual mandatory post-error and post-correct response-to-stimulus times by measuring them based on timestamps on experimental file logs and found that D_err_ = 3136 ms, and D_corr_ = 6370 ms (Fig. S11).

#### Determining D_RSI_

Mean normalized decision time, as described above, is calculated by dividing the mean decision time ⟨*DT*⟩ by D_tot_, equal to T_0_ + D_RSI_, the response-to-stimulus interval [31]. The response-to-stimulus interval comprises two components, the mandatory response-to-stimulus interval (punishment or reward time, times for auditory cues, mandatory intertrial interval, etc.) (Fig. S11), and any extra voluntary intertrial interval (Fig. S12):

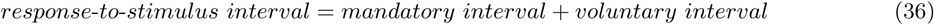

We assume that the animals optimize reward rate based on task engagement time, the sum of reaction times plus all mandatory task time, exiting the task during any extra voluntary inter-trial intervals. Thus, for the purposes of mean normalized decision time:

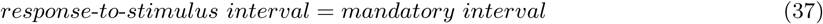

A derivation concluded that the response-to-stimulus interval after an error trial was the only relevant interval (*see* Mean Normalized Decision Time Depends only on T_0_ and D_err_ for derivation), thus:

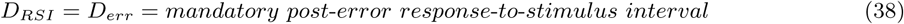

#### Voluntary Intertrial Interval

We conducted a detailed analysis of the voluntary intertrial intervals after both correct and error trials (Fig. S12). To prevent a new trial from initiating while the animals were licking one of the side lickports, the task included a 300 ms interval at the end of a trial where an extra 500 ms were added if the animal licked one of the side lickports (Fig. S11). There was no stimulus (visual or auditory) to indicate the presence of this task feature so the animals were not expected to learn it. It was clear that the animals did not learn this task feature as most voluntary intertrial intervals are clustered in 500 ms intervals and decay after each boundary (Fig. S12a). Aligning the voluntary intertrial distributions every 500 ms reveals substantial overlap (Fig. S12c, d), indicating similar urgency in every 500 ms interval, with an added amount of variance the farther the interval from zero. Moreover, measuring the median voluntary inter-trial interval from 0-500, 0-1000 and 0-2000 ms showed very similar values (47, 67, 108 ms after error trials, Fig. S12b). The median was higher after correct trials (55, 134, 512 ms, Fig. S12b) because the animals were collecting reward from the side lickports and much more likely to trigger the extra 500 ms penalty times.

#### Reward Rate Sensitivity to T_0_ and Voluntary Intertrial Interval

To ensure that our results did not depend on our chosen estimate for T_0_ and our choice to ignore voluntary intertrial intervals when computing metrics like D_*RSI*_ and reward rate, we computed fraction maximum instantaneous reward rate as a function of T_0_ and vountary intertrial interval. We conducted this analysis across n = 26 rats at asymptotic performance (Fig. S13a, b), and during the learning period (Fig. S13c, d). During asymptotic performance, sweeping T_0_ from our estimated minimum to our maximum possible values generated negligible changes in reward rate across a much larger range of possible voluntary intertrial intervals than we observed (Fig. S13a). Reward rate was more sensitive to voluntary intertrial intervals, but did not drop below 90% of the possible maximum when considering a median voluntary intertrial interval up to 2000 ms (the median when allowing up to a 2000 ms window after a trial, after which agents are considered to have “exited the task”) (Fig. S13b). During learning, we found similar results, with possible voluntary intertrial interval values have a larger effect on reward rate than T_0_, however even with the most extreme combination of a maximum T_0_ = 350 ms, and the median voluntary intertrial interval up to 2000ms (Fig. S13d, light grey trace), fraction maximum reward rate was at most 10-15% away from the least extreme combination of T_0_ = 350 ms and voluntary intertrial interval = 0 (Fig. S13c, horizontal line along the bottom of the heat map) for most of the learning period. These results confirm that our qualitative findings do not depend on our estimated values of T_0_ and choice to ignore voluntary intertrial intervals.

#### Ignore Trials

Because of the free-response nature of the task, animals were permitted to ignore trials after having initiated them (Fig. S2). Although the fraction of ignored trials did seem to be higher at the beginning of learning for the first set of stimuli the animals learned (stimulus pair 1; Fig. S2a), this effect did not repeat for the second set (stimulus pair 2, Fig. S2b). This suggests that the cause for ignoring the trials during learning was not stimulus-based but rather related to learning the task for the first time. Overall, the mean fraction of ignored trials remained consistently low across stimulus sets and ignore trials were excluded from our analyses.

#### Post-Error Slowing

In order to verify whether the increase in reaction time we saw at the beginning of learning relative to the end of learning was not solely attributable to a post-error slowing policy, we quantified the amount of post-error slowing during learning for both stimulus pair 1 and stimulus pair 2. For stimulus pair 1, we found that there was a consistent but slight amount of average post-error slowing. (Fig. S7a). This amount was not significantly different at the start and end of learning (Fig. S7b).

We re-did this analysis for stimulus pair 2 and found similar results: animals had a consistent, modest amount of post-error slowing but it did not change across sessions during learning (Fig. S7c). We tested for a significant difference in post-error slowing between the last 2 sessions of stimulus pair 1 and the first 2 sessions of the completely new stimulus pair 2 and found none (Fig. S7d) even though there was a large immediate change in error rate. In fact, there was a trend towards a decrease in post-error slowing (and towards post-correct slowing) in the first few sessions of stimulus pair 2. This is consistent with the hypothesis that post-error slowing is an instance of a more general policy of orienting towards infrequent events [54]. As correct trials became more infrequent than error trials when stimulus pair 2 was presented, we observed a trend towards post-correct slowing, as predicted by this interpretation.

Our subjects exhibit a modest, consistent amount of post-error slowing, which could at least partially explain the reaction time differences we see throughout learning. An experiment with transparent stimuli where error rate was constant but reaction times dropped, however, strongly contradicts the account that the rats implement a simple strategy like post-error slowing to modulate their reaction times during learning.

### Recurrent Neural Network Model and Learning DDM (LDDM) Reduction

We consider a recurrent network receiving noisy visual inputs over time. In particular, we imagine that an input layer projects through weighted connections to a single recurrently connected read-out node, and that the weights must be tuned to extract relevant signals in the input. The read-out node activity is compared to a modifiable threshold which governs when a decision terminates. This network model can then be trained via error-corrective gradient descent learning or some other procedure. In the following we derive the average dynamics of learning.

To reduce this network to a drift-diffusion model with time-dependent SNR, we first note that due to the law of large numbers, activity increments of the read-out node will be Gaussian provided that the distribution of input stimuli has bounded moments. We can thus model the input-to-readout pathway at each time step as a Gaussian input *x*(*t*) flowing through a scalar weight *u*, with noise of variance 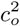 added before the signal is sent into an integrating network. Taking the continuum limit, this yields a drift-diffusion process with effective drift rate *Ã* = *Au* and noise variance 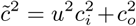. Here *A* parameterizes the perceptual signal, 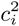 is the input noise variance (noise in input channels that cannot be rejected), and 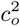 is the output noise variance (internal noise in output circuitry). The resulting decision variable *ŷ* at time *T* is Gaussian distributed as 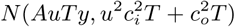 where *y* is the correct binary choice. A decision is made when *ŷ* hits a threshold of ±*z*.

### Within-trial Drift-diffusion Dynamics

On every trial, therefore, the subject’s behavior is described by a drift-diffusion process, for which the average reward rate as a function of signal to noise and threshold parameters is known [10]. The accuracy and decision time of this scheme is determined by two quantities. First, the signal-to-noise ratio

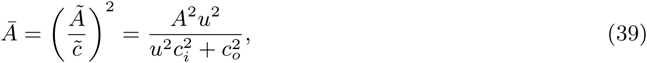

and second, the threshold-to-drift ratio 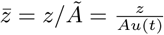. We can rewrite the signal-to-noise ratio as

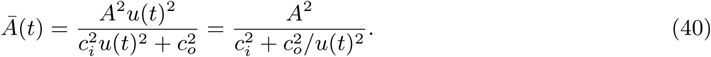

From this it is clear that, when learning has managed to amplify the input signals such that *u*(*t*) → ∞, the asymptotic signal-to-noise ratio is simply 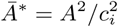. Further, rearranging to

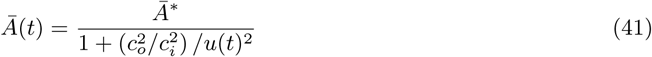

shows that there are in fact just two parameters: the asymptotic achievable SNR *Ā*^*^ and the output-to input-noise variance ratio 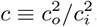,

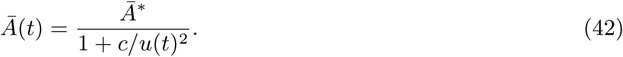

The mean error rate (ER), mean decision time (DT), and mean reward rate (RR) are therefore

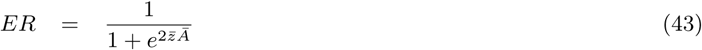

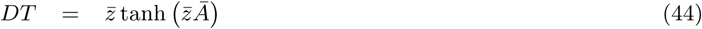

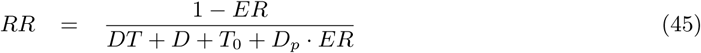

where we have suppressed the dependence of *Ā* and 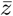 on time for clarity. Here *D* is the interval between a correct response and the next trial, *T*_0_ is the time required for non-decision making processing (e.g., motor responses or initial sensory delays), and *D*_*p*_ is extra penalty time added to error responses.

The term 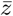*Ā* is a measure of the total evidence accrued on average, and is equal to

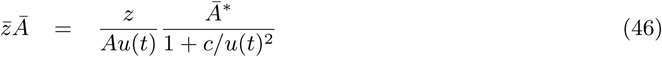

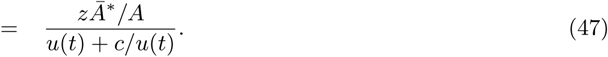

Here for a fixed threshold *z*, the denominator shows the trade-off for increasing perceptual sensitivity: small *u*(*t*) causes errors due to output noise, while large *u*(*t*) causes errors due to overly fast integration for the specified threshold level.

#### Across-trial Error-Corrective Learning Dynamics

To model learning, we consider that animals adjust perceptual sensitivities *u* over time in service of minimizing an objective function. In this section we derive the average learning dynamics when the objective is to minimize the error rate. The Learning DDM (LDDM) can be conceptualized as an “outer-loop” that modifies the SNR of a standard DDM “inner-loop” described in the preceding subsection. If perceptual learning is slow, there is a strong separation of timescales between these two loops. On the timescale of a single trial, the agent’s SNR is approximately constant and evidence accumulation follows a standard DDM, whereas on the timescale of many trials, the specific outcome on any one trial has only a small effect on the network weights *w*, such that the learning-induced changes are driven by the *mean* ER and DT.

To derive the mean effect of error-corrective learning updates, we suppose that on each trial the network uses gradient descent on the hinge loss to update its parameters, corresponding to standard practice for supervised neural networks. The hinge loss is

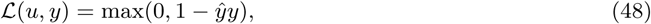

yielding the gradient descent update

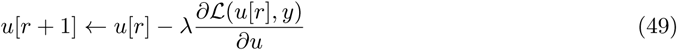

where *λ* is the learning rate and *r* is the trial number.

When the learning rate is small (*λ* ≪ 1), each trial changes the weights minimally and the overall update is approximately given by the average continuous-time dynamics

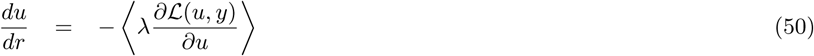

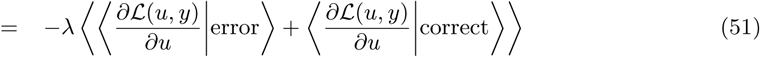

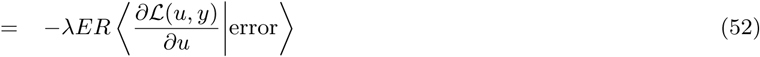

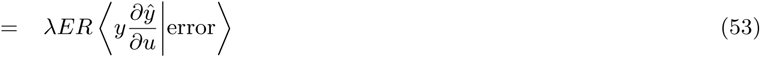

where ⟨·⟩ denotes an average over the correct answer *y*, the inputs and the output noise. The first step follows from iterated expectation. The second step follows from the fact that the probability of an error is simply the error rate *ER*, and for correct trials, the derivative of the hinge loss is zero. Next,

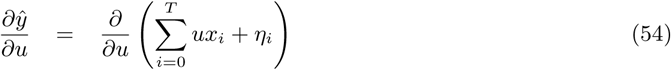

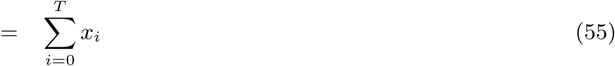

where *T* is the time step at which *ŷ* crosses the decision threshold ±*z*. Returning to Eq. (53),

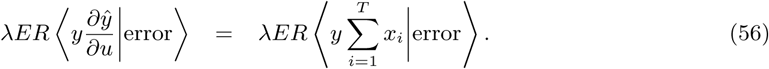

Hence the magnitude of the update depends on the typical total sensory evidence given that an error is made. To calculate this, let 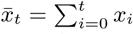 be the total sensory evidence up to time *t*, and 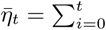 be the total decision noise up to *t*. These are independent and normally distributed as

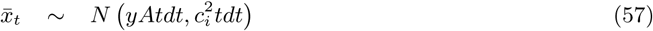

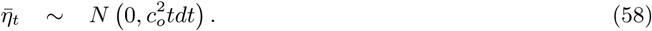

Therefore, we have

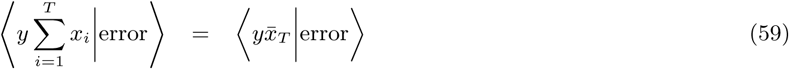

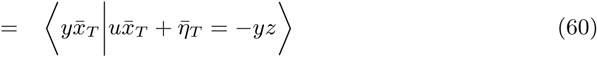

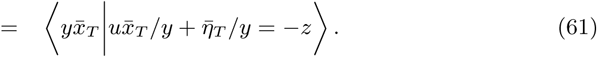

These variables are jointly Gaussian. Letting 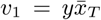 and 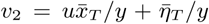, the means *µ*_1_, *µ*_2_, variances 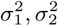, and covariance Cov(*v*_1_, *v*_2_) of *v*_1_, *v*_2_ given the hitting time *T* are

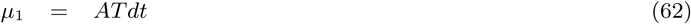

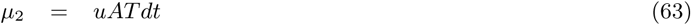

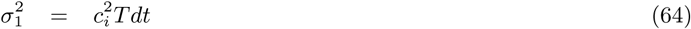

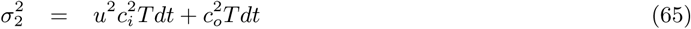

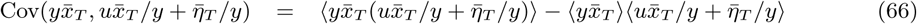

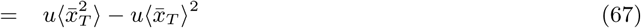

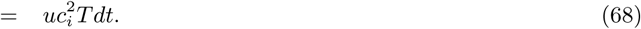

The conditional expectation is therefore

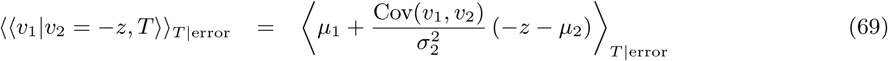

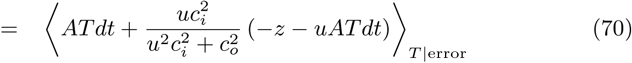

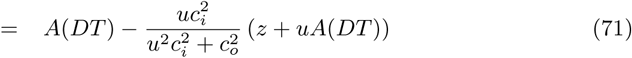

where we have used the fact that ⟨*T dt*⟩ _*T* | error_ = *DT*, because in the DDM model the mean decision time is the same for correct and error trials. Inserting Eq. (71) into Eq. (56) yields

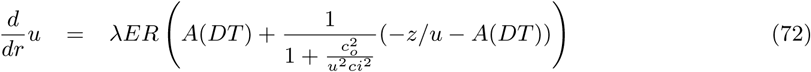

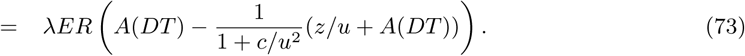

Finally, we switch the units of the time variable from trials to seconds using the relation *dt* = *D*_*tot*_*dr*, yielding the dynamics

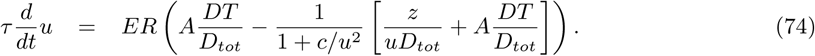

The above equation describes the dynamics of *u* under gradient descent learning. We note that here, the dependence of the dynamics on threshold trajectory is contained implicitly in the *DT, ER*, and *D*_*tot*_ terms.

To obtain equivalent dynamics for the SNR *Ā*, we have

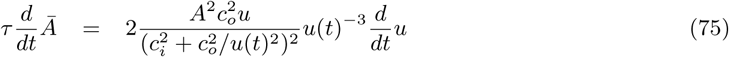

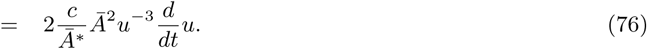

Rearranging the definition of *Ā* yields

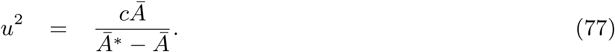

Inserting Eq. (77) into Eq. 76 and simplifying, we have

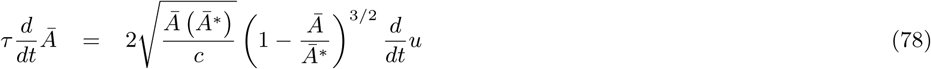

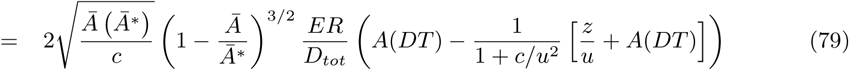

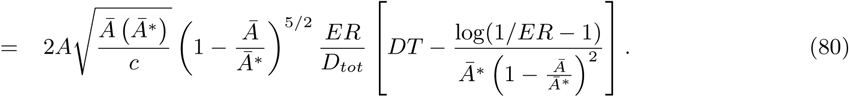

Here in the second step we have used the fact that 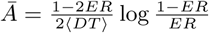 and Eq. (77). Finally, absorbing the drift rate *A* into the time constant 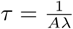, we have the dynamics

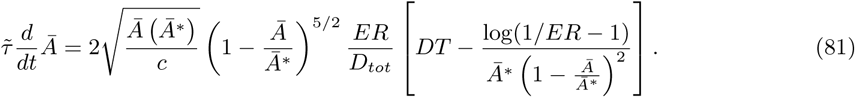

This equation reveals that the Learning DDM has four scalar parameters: the asymptotic SNR *Ā*^*^, the output-to-input-noise variance ratio *c*, the initial SNR at time zero *Ā*(0), and the combined drift-rate/learning rate time constant 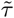. In addition, it requires the choice of threshold trajectory *z*(*t*).

To reveal the basic learning speed/instantaneous reward rate trade-off in this model, we investigate the limit where *Ā* is small but finite (low signal-to-noise) and the threshold is small, such that the error rate is near *ER* = 1*/*2. Then the second term in Eq. (81) goes to zero, giving

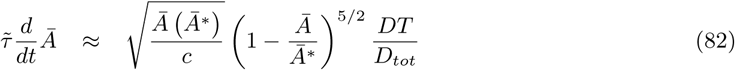

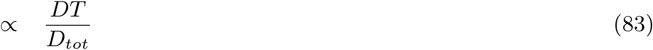

such that learning speed is increasing in *DT*. By contrast the instantaneous reward rate when *ER* = 1*/*2 is

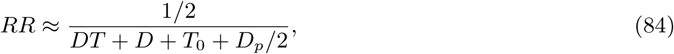

which is a decreasing function of *DT*.

#### Threshold Policies

We evaluate several simple threshold policies. The iRR-greedy policy sets 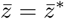, the instantaneous reward rate optimal policy at all times. The constant threshold policy sets 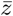 to a fixed constant throughout learning. The iRR-sensitive policy initializes the threshold to a fixed initial condition, and then moves towards the iRR-optimal decision time using the dynamics

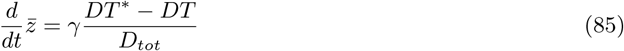

where *γ* controls the rate of convergence.

Finally, the global optimal policy optimizes the entire function 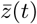 to maximize total cumulative reward during exposure to the task. To compute the optimal threshold trajectory, we discretize the reduction dynamics in Eq.(74) and perform gradient ascent on 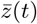 using automatic differentiation in the PyTorch python package. While this procedure is not guaranteed to find the global optimum (due to potential nonconvexity of the optimization problem), in practice we found highly reliable results from a range of initial conditions and believe that the identified threshold trajectory is near the global optimum.

#### Parameter Fitting

The LDDM model has several parameters governing its performance, including the asymptotic optimal SNR, the output/input noise variance ratio, the learning rate, and parameters controlling threshold policies where applicable. To fit these, we discretized the reduction dynamics and performed gradient ascent on the log likelihood of the observed data under the LDDM model, again using automatic differentiation in the Pytorch python package. Because our model is highly simplified, our goal was only to place the parameters in a reasonable regime rather than obtain quantitative fits. We note that our fitting procedure could become stuck in local minima, and that a range of other parameter settings might also be consistent with the data. The best-fitting parameters we obtained and used in all model results were *A* = 0.9542, *c*_*i*_ = 0.3216, *c*_*o*_ = 30, *u*_0_ = .0001. We used a discretization timestep of *dt* = 160. For the constant threshold and iRR-sensitive policies, the best fitting initial threshold was *z*(0) = 30. For the iRR-sensitive policy, the best fitting decay rate was *γ* = 0.00011891.

## Supplementary Information

**Table S1:** Individual Animal Participation Across Behavioral Experiments

| Animal | Sex | Stimulus Pair 1 | Stimulus Pair 2 | Transparent Stimuli | Stimulus Pair 3 |
| --- | --- | --- | --- | --- | --- |
| AK1 | F | size & rotation |  | alpha = 0 |  |
| AK2 | F | size & rotation |  | alpha = 0.1 |  |
| AK3 | F | size & rotation |  | alpha = 0.0 |  |
| AK4 | F | size & rotation |  |  |  |
| AK5 | F | size & rotation |  | alpha = 0.1 (excluded)‡ |  |
| AK6 | F | size & rotation |  | alpha = 0 |  |
| AK7 | F | size & rotation |  | alpha = 0 |  |
| AK8 | F | size & rotation (excluded)* |  |  |  |
| AK9 | F | size & rotation |  | alpha = 0.1 |  |
| AK10 | F | size & rotation |  | alpha = 0.1 |  |
| AK11 | F | size & rotation |  | alpha = 0.0 |  |
| AK12 | F | size & rotation |  | alpha = 0.1 (excluded)‡ |  |
| AL1 | F | size & rotation | canonical only |  | (excluded)§ |
| AL2 | F | size & rotation | canonical only |  | below |
| AL3 | F | size & rotation | canonical only |  | above |
| AL4 | F | size & rotation | canonical only |  | below (excluded)¶ |
| AL5 | F | size & rotation | canonical only |  | below |
| AL6 | F | size & rotation | canonical only |  | below |
| AL7 | F | size & rotation | canonical only |  | below (excluded)¶ |
| AL8 | F | size & rotation | canonical only |  | above |
| AL9 | F | size & rotation |  |  |  |
| AL10 | F | size & rotation |  |  |  |
| AL11 | F | size & rotation |  |  |  |
| AL12 | F | size & rotation (excluded)* |  |  |  |
| AL13 | F | size & rotation | canonical only |  | above |
| AL14 | F | size & rotation | canonical only |  | below |
| AL15 | F | size & rotation | canonical only |  | above |
| AL16 | F | size & rotation | canonical only |  | above |
| AM1 | F | size & rotation† | canonical only |  | below (excluded)\|\| |
| AM2 | F | size & rotation† | canonical only |  | below |
| AM3 | F | size & rotation† | canonical only |  | above |
| AM4 | F | size & rotation† | canonical only |  | above |
| AM5 | F | size & rotation† |  |  |  |
| AM6 | F | size & rotation† |  |  |  |
| AM7 | F | size & rotation† |  |  |  |
| AM8 | F | size & rotation† |  |  |  |
| AN1 | F | canonical only |  |  |  |
| AN2 | F | canonical only |  |  |  |
| AN3 | F | canonical only |  |  |  |
| AN4 | F | canonical only |  |  |  |
| AN5 | F | canonical only |  |  |  |
| AN6 | F | canonical only |  |  |  |
| AN7 | F | canonical only |  |  |  |
| AN8 | F | canonical only |  |  |  |

Exclusions \* failed to learn task. † not included in initial learning experiment. ‡ above chance for near-transparent stimuli. § failed to learn previous stimuli. ¶ not enough practice trials with reaction time restrictions. || failed to learn stimuli with reaction time restrictions.

**Figure S1:**
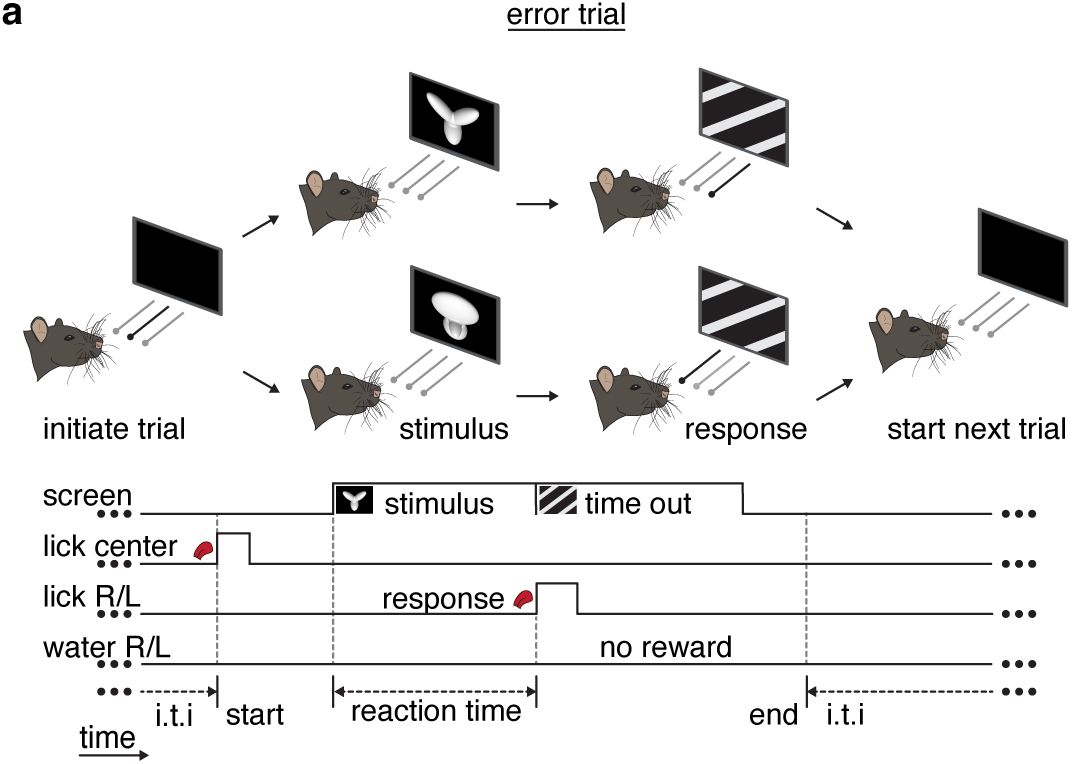
Task schematic for error trials. **(a)** Error trial: rat chooses incorrect left/right response port and incurs a timeout period.

**Figure S2:**
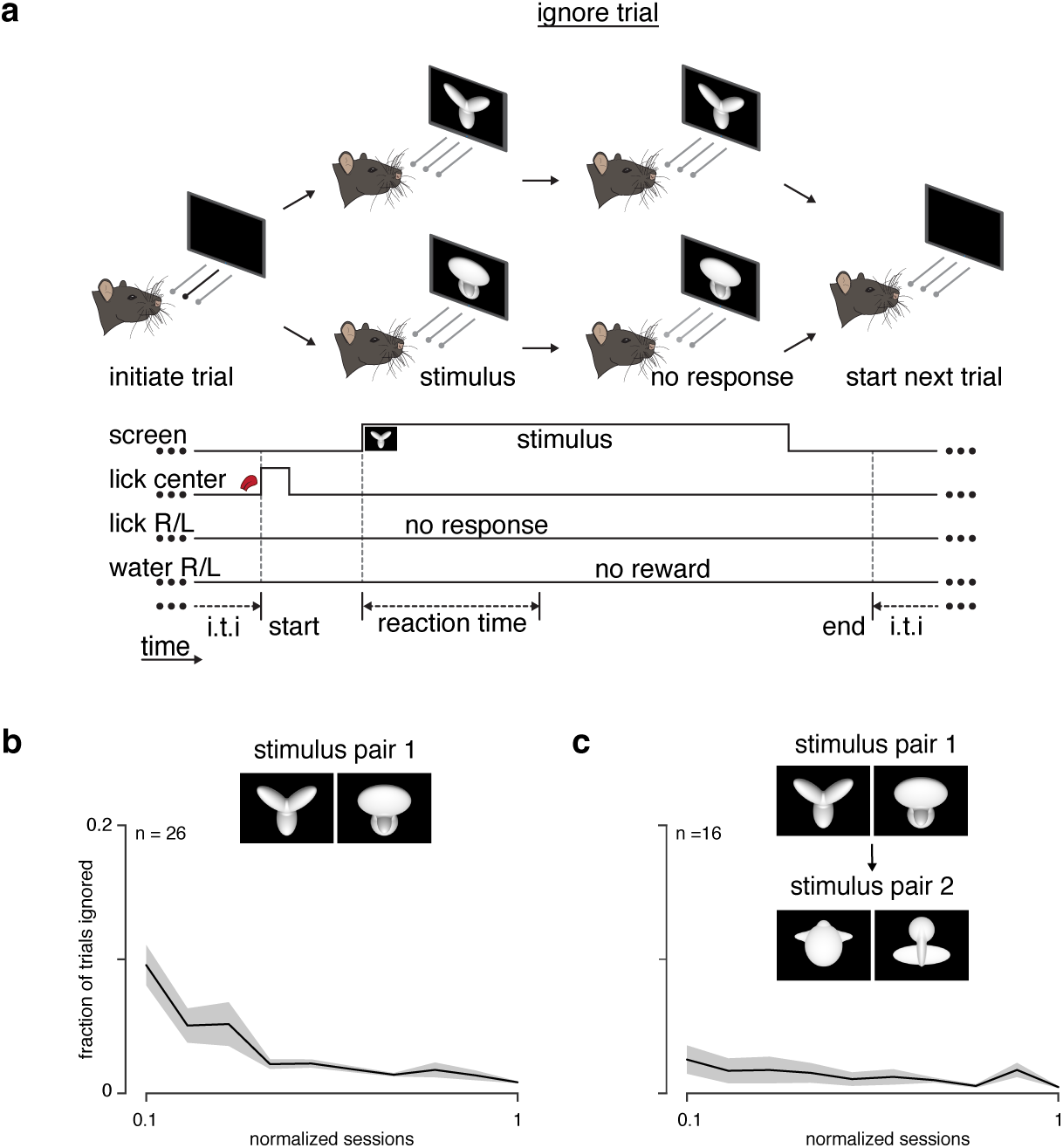
Fraction of ignored trials during learning. **(a)** Schematic of an ignore trial: rat does not choose a left/right response port and receives no feedback. **(b)** Fraction of trials ignored (ignored trials / (correct + incorrect + ignored trials)) during learning for animals encountering the task for the first time (stimulus pair 1). **(c)** Fraction of trials ignored for animals learning stimulus pair 2 after training on stimulus pair 1.

**Figure S3:**
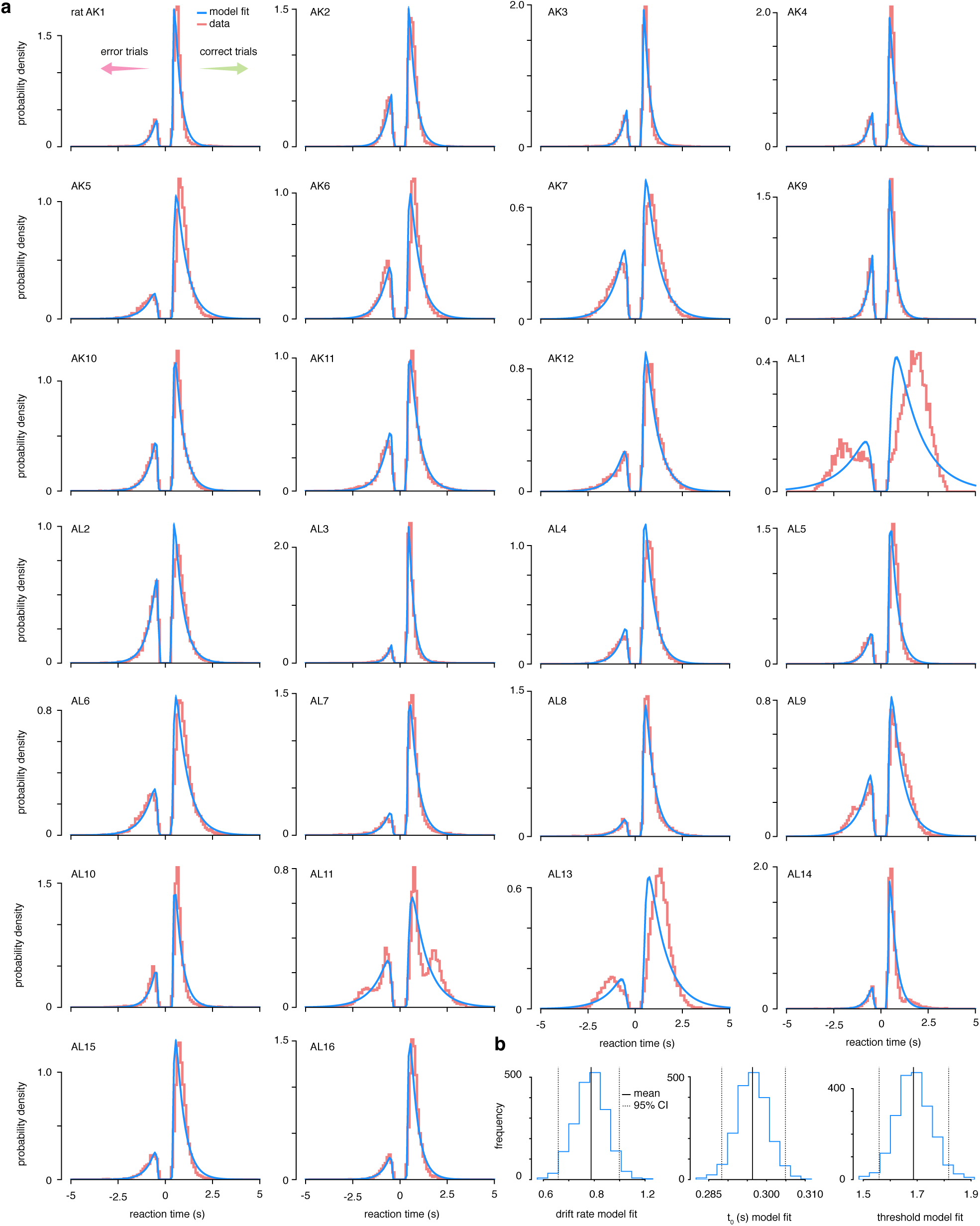
Drift-diffusion model data fits. **(a)** The accuracy and reaction time data from 26 trained rats was fit to a simple drift-diffusion model using the hierarchical Bayesian estimation of the drift-diffusion model (HDDM) package [107] **(b)** Estimated parameter value across all animals. The parameter estimates are distributions because of the Bayesian nature of the estimation method.

**Figure S4:**
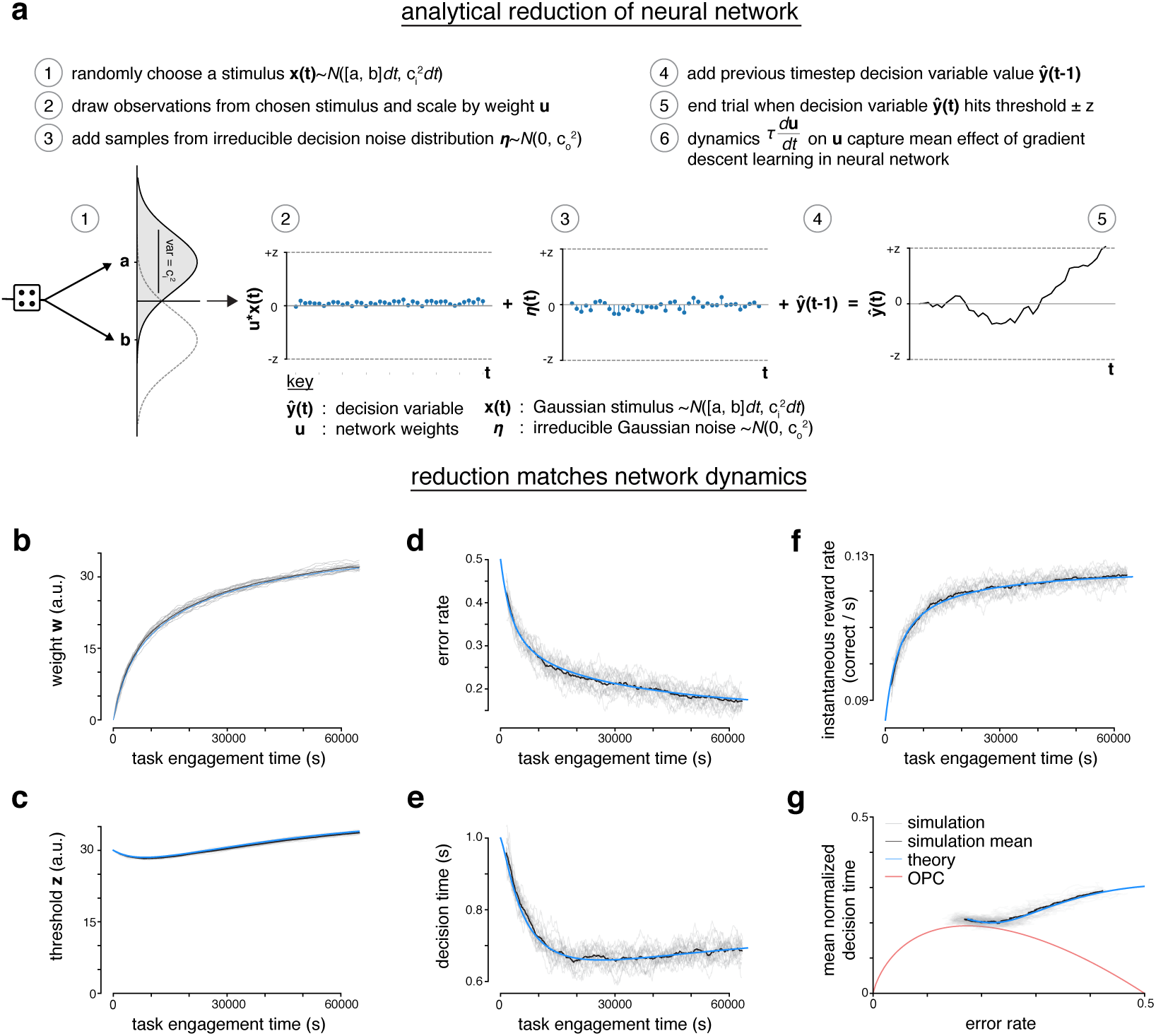
Analytical reduction of LDDM matches error-corrective learning neural network dynamics during learning. **(a)** The recurrent linear neural network can be analytically reduced. In the reduction, the decision variable draws an observation from one of two randomly chosen Gaussian “stimuli.” The observations are scaled by a perceptual weight. After the addition of some irreducible noise, the value of the decision variable at previous time step is added to the current time step. A trial ends once the decision variable hits a predetermined threshold. The dynamics of the perceptual weight capture the mean effect of gradient descent learning in the recurrent linear neural network. **(b)** Weight **w** of neural network across task engagement time for multiple simulations of the network (grey), the mean of the simulations (black) and the analytical reduction of the network (blue). **(c)** Same as in **b** but for the threshold **z. (d)** Same as in **b** but for the error rate **(e)** Same as in **b** but for the decision time **(f)** Same as in **b** but for the instantaneous reward rate (correct trials per second) **(g)** Learning trajectory in speed-accuracy space for simulations, simulation mean and analytical reduction (theory). OPC is shown in red.

**Figure S5:**
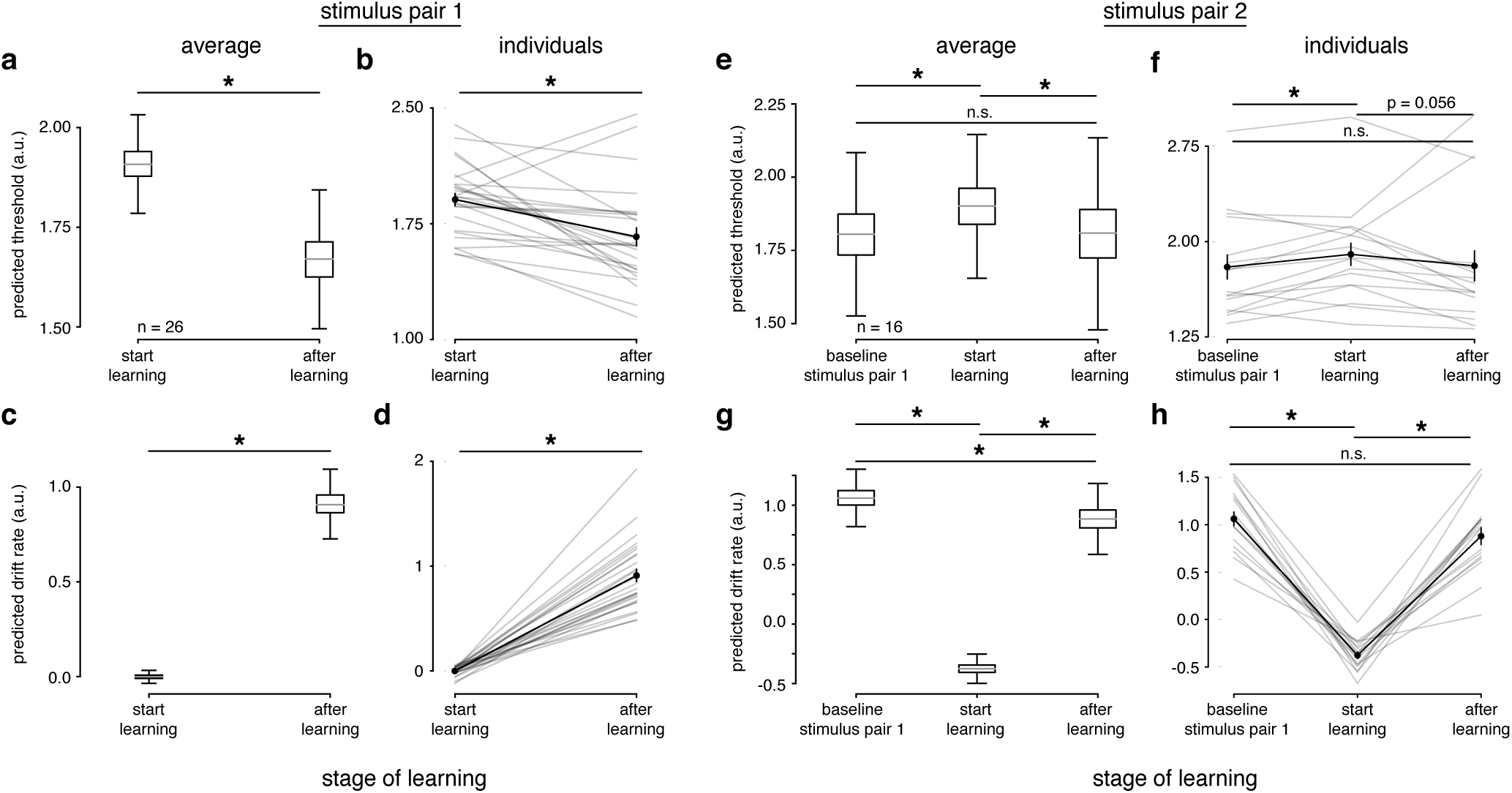
Simple DDM fits indicate threshold decreases and drift rate increases during learning. **(a)** The learning data from stimulus pair 1 and 2 were fit with a simple DDM using the HDDM framework [107]. The HDDM framework reports a distribution of possible parameter values for the population (**a, c, e, g**), as well as fits for individuals (**b, d, f, h**). **(a)** Predicted threshold values for average model across rats (n = 26) for start of learning and after learning (first 1000 and last 1000 trials for each subject); ‘start’ and ‘after’ (p ¡ 1e-4). **(b)** Same as **a** but for individuals (grey and their mean and SEM (black); ‘start’ and ‘after’ learning (p = 0.0006). **(c)** Predicted drift rate values for average model across rats (n = 26) for start of learning and after learning (p ¡ 1e-4). **(d)** Same as **c** but for individuals (grey) and their mean and SEM (black); ‘start’ and ‘after’ (p ¡ 1e-4). (e) Predicted threshold values for average model across rats (n = 16) for baseline trials with stimulus pair 1, start of learning and after learning of stimulus pair 2 (last 500 trials with stimulus pair 1, first 500 and last 500 trials with stimulus pair 2 for each subject); baseline and ‘start’ (p = 0.0386), ‘start’ and ‘after’ (p = 0.0557), baseline and ‘after’ (p = 0.8767). **(f)** Same as **e** but for individuals (grey and their mean and SEM (black); baseline and ‘start’ (p ¡ 1e-4), ‘start’ and ‘after’ (p ¡ 1e-4), baseline and ‘after’ (p = 0.301601). **(g)** Predicted drift rate values for average model across rats (n = 16) for baseline, start of learning and after learning; baseline and ‘start’ (p ¡ 1e-4), ‘start’ and ‘after’ (p ¡ 1e-4), baseline and ‘after’ (p ¡ 1e-4). **(h)** Same as **g** but for individuals (grey) and their mean and SEM (black); baseline and ‘start’ (p = 0.0004), ‘start’ and ‘after’ (p = 0.0004), baseline and ‘after’ (p = 0.1089). Wilcoxon signed-rank test for all tests, * denotes p ¡ 0.05, n.s. denotes ‘not significant’ (p ¿ 0.05).

**Figure S6:**
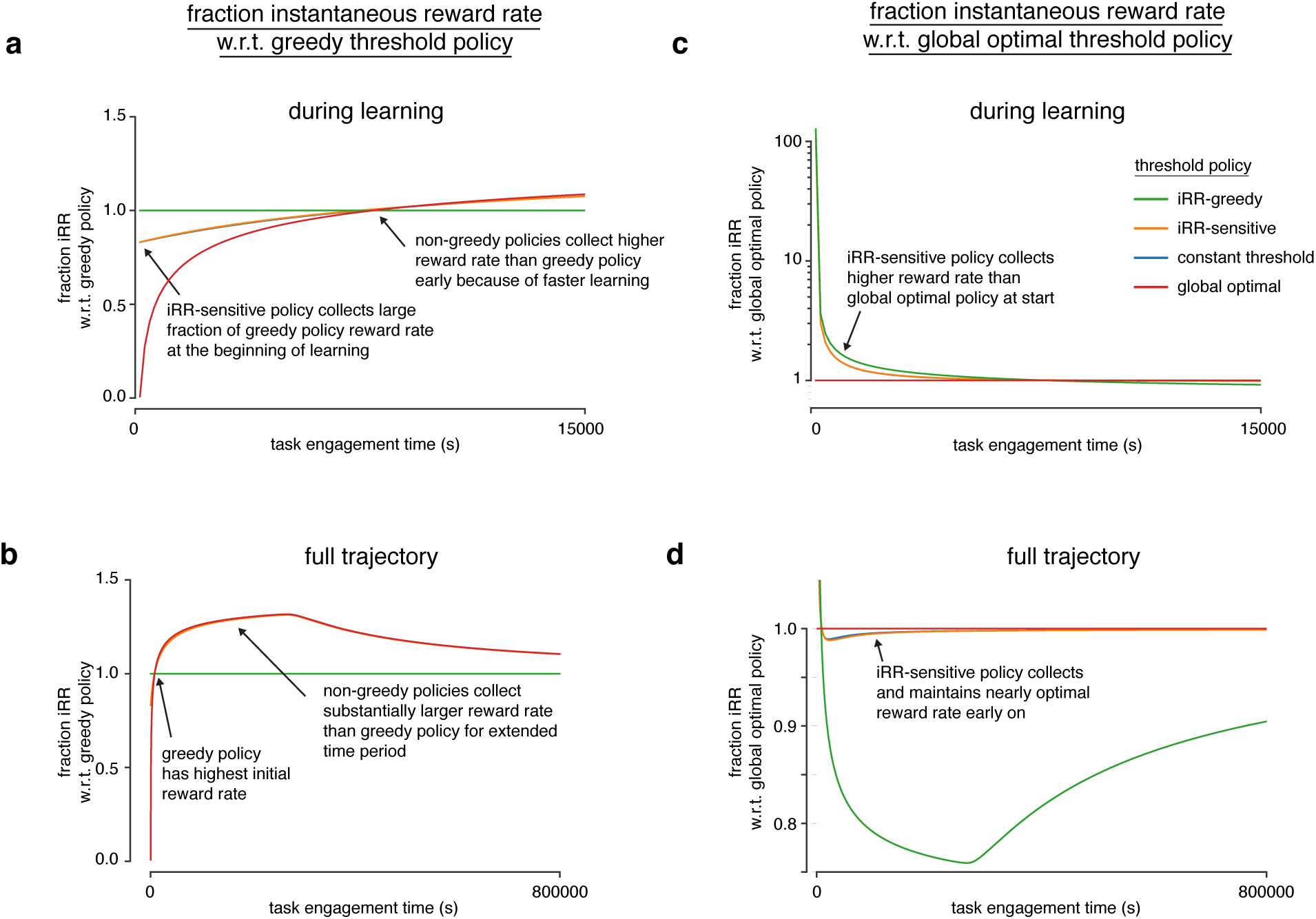
Model reveals rat learning dynamics resemble optimal trajectory without relinquishing initial rewards. **(a)** Fraction of instantaneous reward rate with respect to the iRR-greedy policy for all model threshold policies during learning. The instantaneous reward rates of all policies were normalized by the iRR-greedy policy’s instantaneous reward rate through task engagement time. **(b)** Same as **a** but for the full trajectory of the simulation. **c** Fraction of instantaneous reward rate with respect to the global optimal policy for all model threshold policies during learning. The instantaneous reward rates of all policies were normalized by the greedy policy’s instantaneous reward rate through task engagement time. **(d)** Same as **c** but for the full trajectory of the simulation.

**Figure S7:**
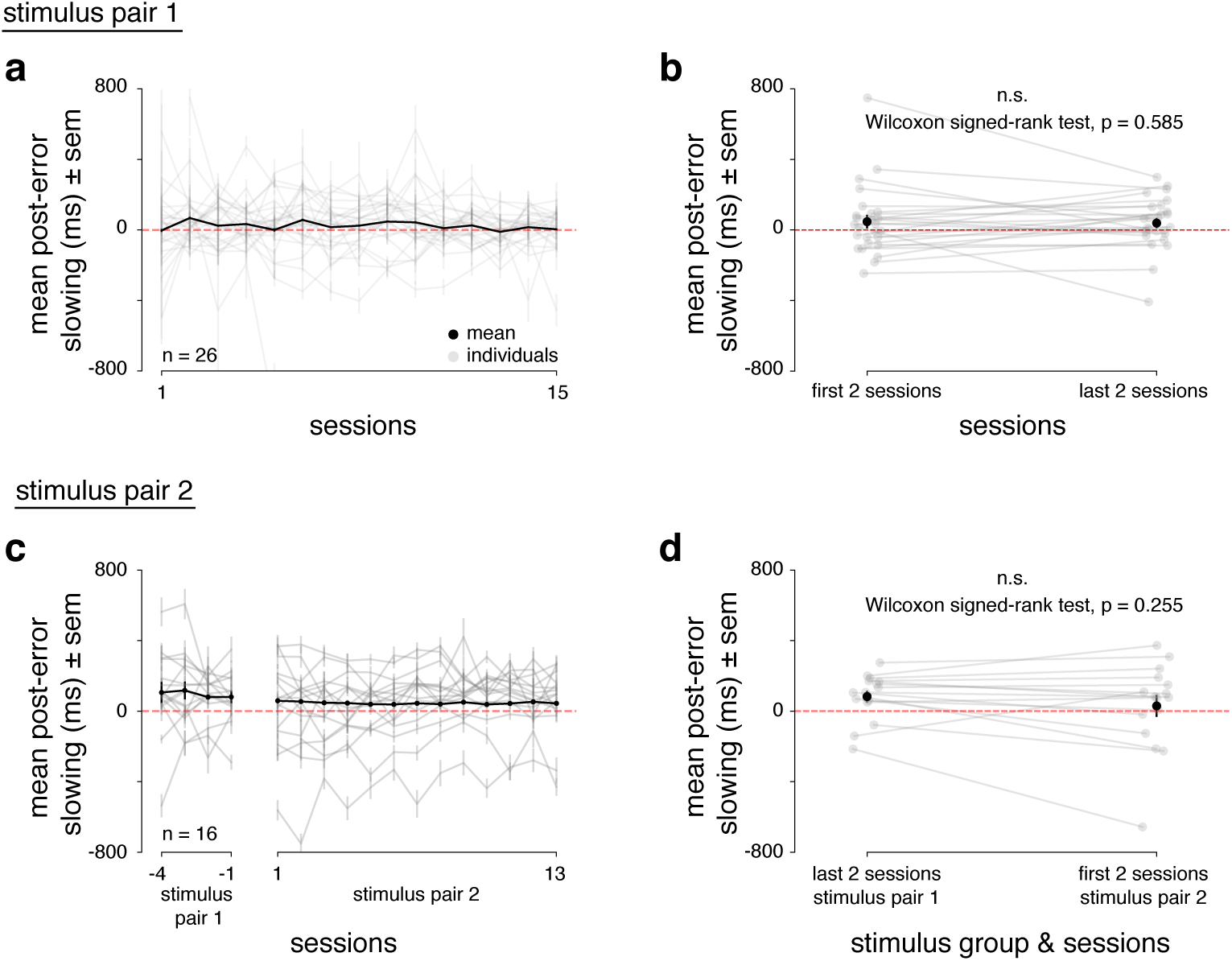
Post-error slowing during rat learning dynamics. **(a)** Individual (grey) and mean (black) post-error slowing across first 15 sessions for n = 26 animals. Post-error slowing was calculated by taking the difference between RTs on trials with previous correct trials and previous error trials. A positive difference indicates post-error slowing. **(b)** Individual mean (grey) and population mean (black) post-error slowing for first 2 sessions of learning and last 2 sessions of learning for n = 26 animals. A Wilcoxon signed-rank test found no significant difference in post-error slowing between the first 2 and last 2 sessions for every animal (p = 0.585). **(c)** Same as in **a** for n = 16 rats, with the addition of 4 baseline sessions with stimulus pair 1 plus the 13 sessions while subjects were learning stimulus pair 2. **(d)** Same as in **b** but comparing the last 2 baseline sessions with stimulus pair 1 and the first 2 sessions learning stimulus pair 2. A Wilcoxon signed-rank test found no significant difference in post-error slowing when the animals started learning stimulus pair 2 (p = 0.255).

**Figure S8:**
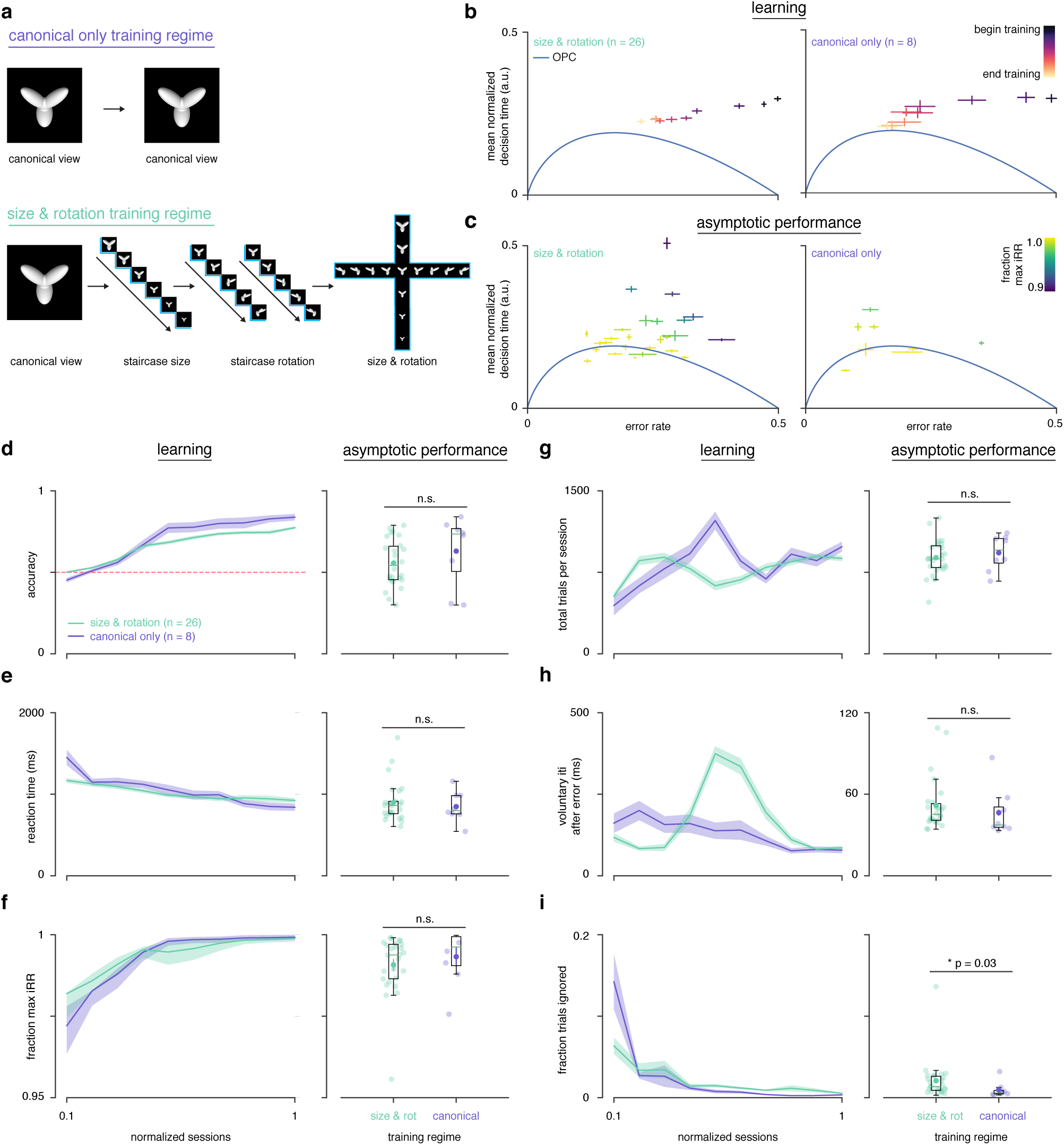
Comparison of training regimes. **(a)** ‘Canonical only’: rats trained to asymptotic performance with only front-view image of each of the two stimuli. ‘Size & rotation’: rats first shown front-view image of stimuli. After reaching criterion (accuracy = 0.7), size staircased. Following criterion, rotation staircased. Upon criterion, stimuli randomly drawn across size and rotation. **(b)** Learning trajectory in speed-accuracy space over normalized training time for rats trained with the ‘size & rotation’ (left panel) and the ‘canonical only’ training regimes (right panel). **(c)** Average location in speed-accuracy space for 10 sessions after asymptotic performance for individual rats in both training regimes, as in **b. (d)** Mean accuracy over learning (left panel) and for 5 sessions after asymptotic performance (right panel) for rats trained with the ‘size & rotation’ (n = 26) and the ‘canonical only’ (n = 8) training regimes. **(e)** Mean reaction time. **(f)** Mean fraction max iRR. **(g)** Mean total trials per session. **(h)** Mean voluntary intertrial interval up to 500 ms after error trials. **(i)** Mean fraction ignored trials. All errors are SEM. Significance in right panels of **d-i** determined by Wilcoxon rank-sum test with p ¡ 0.05.

**Figure S9:**
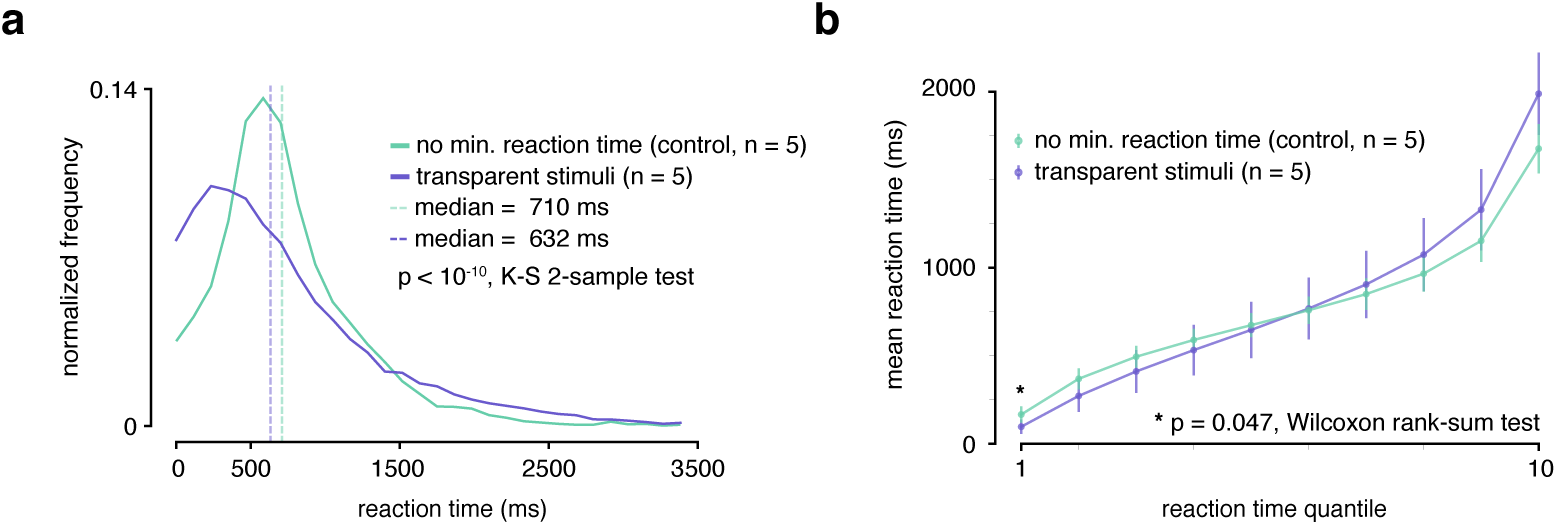
Reaction time analysis of transparent stimuli experiment. **(a)** During transparent stimuli, the reaction time (RT) minimum was relaxed to 0 ms to fully measure a possible shift in RT behavior. To be able to ascertain whether transparent stimuli led to a significant chance in RT, the RT histogram of transparent stimuli (purple) sessions was compared to control sessions with visible stimuli (green) with no RT minimum. Medians indicated with dashed lines. Kolmogorv-Smirnov 2-sample test over distributions found significant different (p¡10^−^10 **(b)** Vincentized RTs for transparent and control visible stimuli sessions with no minimum reaction time. A Mann-Whitney U Test for the first quantile (fastest RTs) found a significant difference between the two groups (p = 0.047).

**Figure S10:**
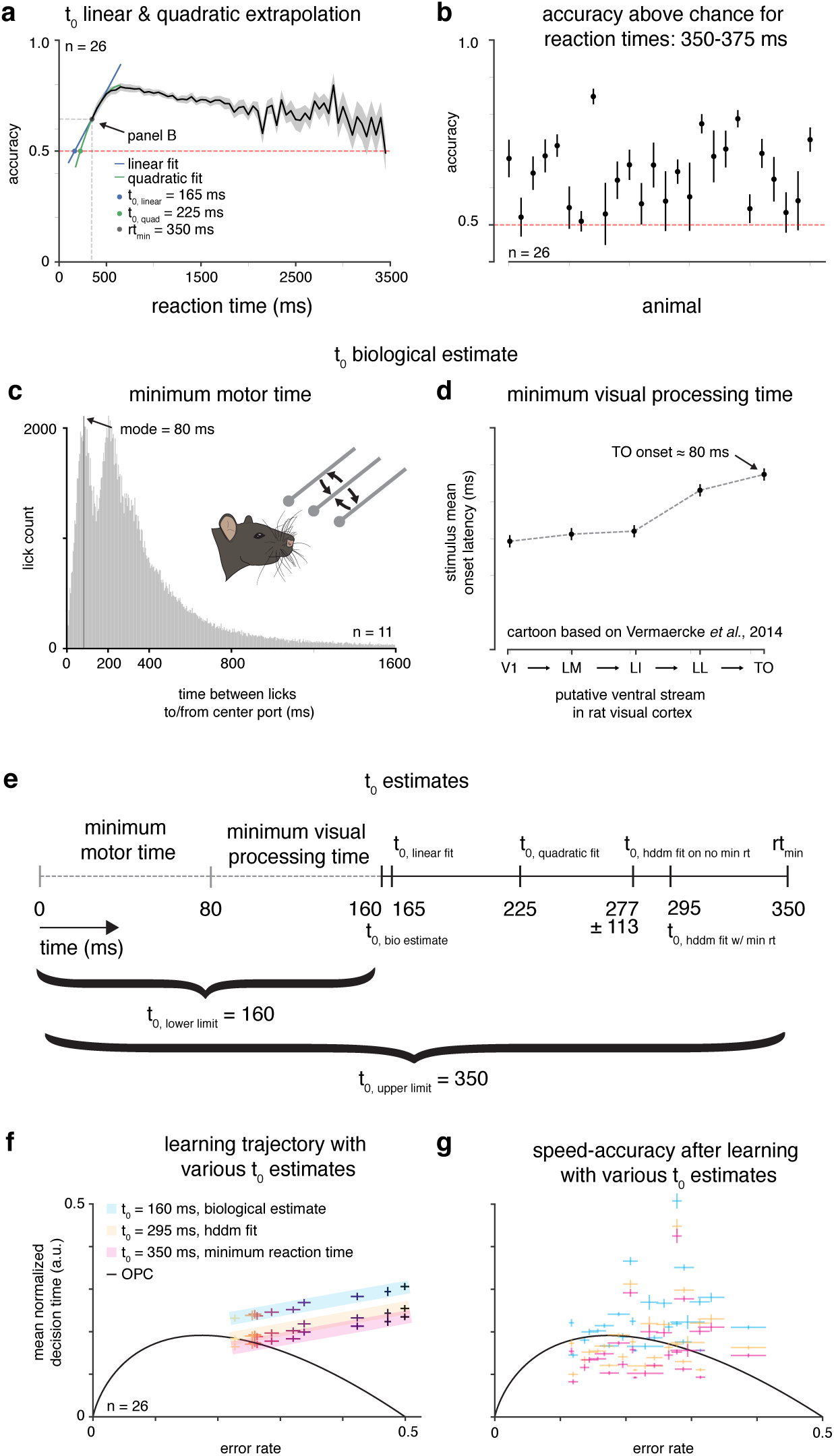
Estimating T_0_. **(a)** Linear and quadratic extrapolations to accuracy as a function of reaction time. The t_0_ estimate is when each extrapolation intersects chance accuracy (0.5). **(b)** Mean accuracy for trials with reaction times 350-375 ms for n = 26 rats. **(c)** Minimum motor time estimated by looking at first peak of time between licks to/from center port for n = 11 rats. **(d)** Cartoon of stimulus onset latency across visual areas from Vermaercke et al., 2014 [112] to estimate minimum visual processing time. **(e)** Diagram of t_0_ estimates, with an upper limit (minimum reaction time) and lower limit (minimum motor time + minimum visual processing time). **(f)** Mean learning trajectory for n = 26 rats with various t_0_ estimates. **(g)** Subjects (n = 26) in speed-accuracy space with various t_0_ estimates.

**Figure S11:**
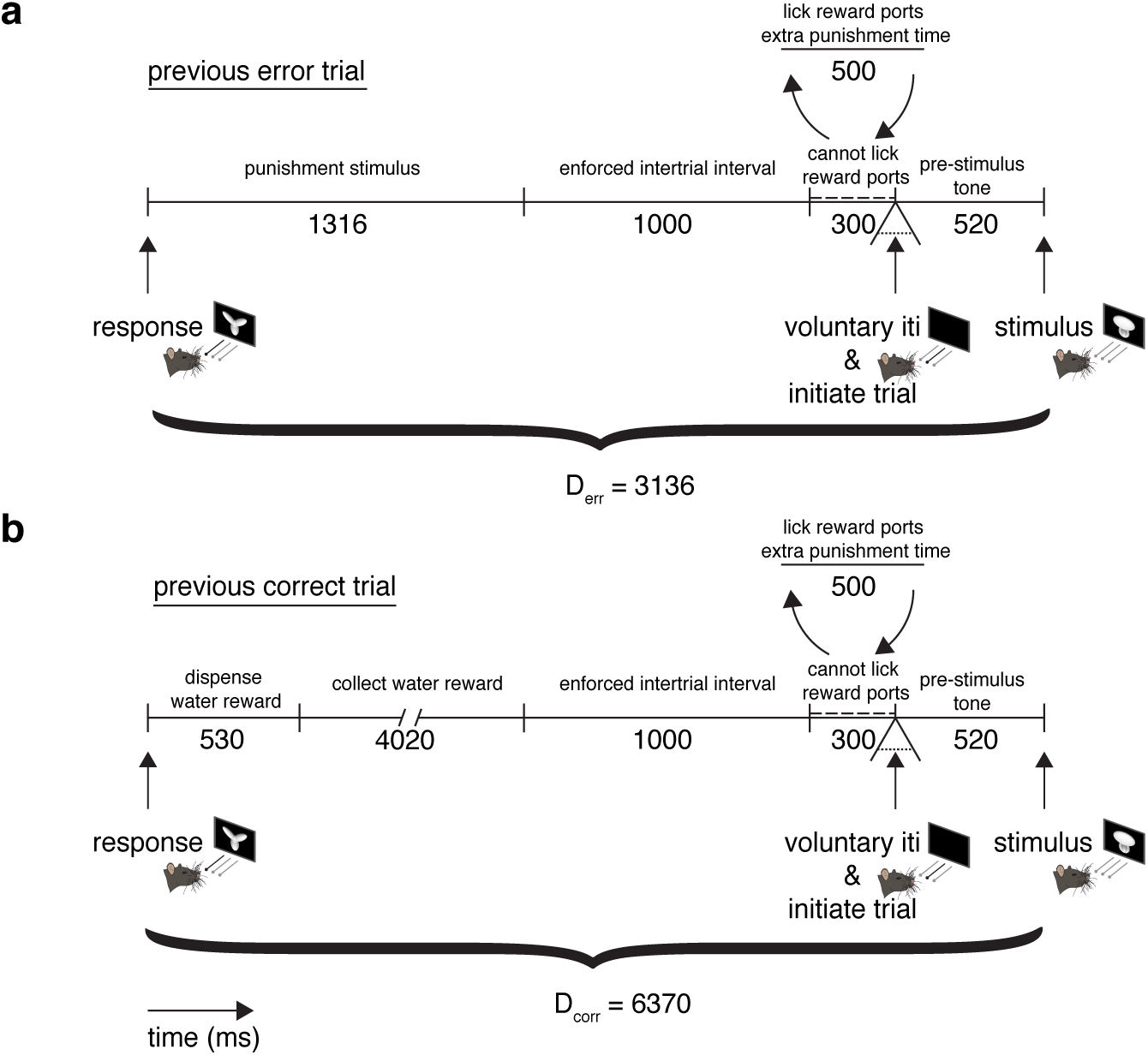
Mandatory post-error (Derr) and post-correct (Dcorr) response-to-stimulus interval times. **(a)** Diagram of intertrial interval (ITI) after previous error trial. All times (punishment stimulus, enforced intertrial interval, cannot lick reward ports and pre-stimulus time) were verified based on timestamps on experimental file logs. After the punishment stimulus and enforced intertrial interval, there is a 300 ms period where rats cannot lick the reward ports. If violated, 500 ms are added to the intertrial interval followed by another 300 ms ‘cannot lick’ period. In addition to this restriction, rats may take as much voluntary time between trials as they wish. Any violation of the ‘cannot lick’ period is counted as voluntary time, and only the minimum mandatory time of 3136 ms is counted for D_err_. **(b)** Diagram of ITI after previous correct trial. All times (dispense water reward, collect water reward, enforced intertrial interval, cannot lick reward ports, pre-stimulus time) was verified based on timestamps on experimental file logs. The same ‘cannot lick’ period is present as in **a**. All times (dispense water reward, collect water reward, enforced intertrial interval, cannot lick reward ports, pre-stimulus time) was verified based on timestamps on experimental file logs. Any violation of the ‘cannot lick’ period is counted as voluntary time, and only the minimum mandatory time of 6370 ms is counted for D_corr_.

**Figure S12:**
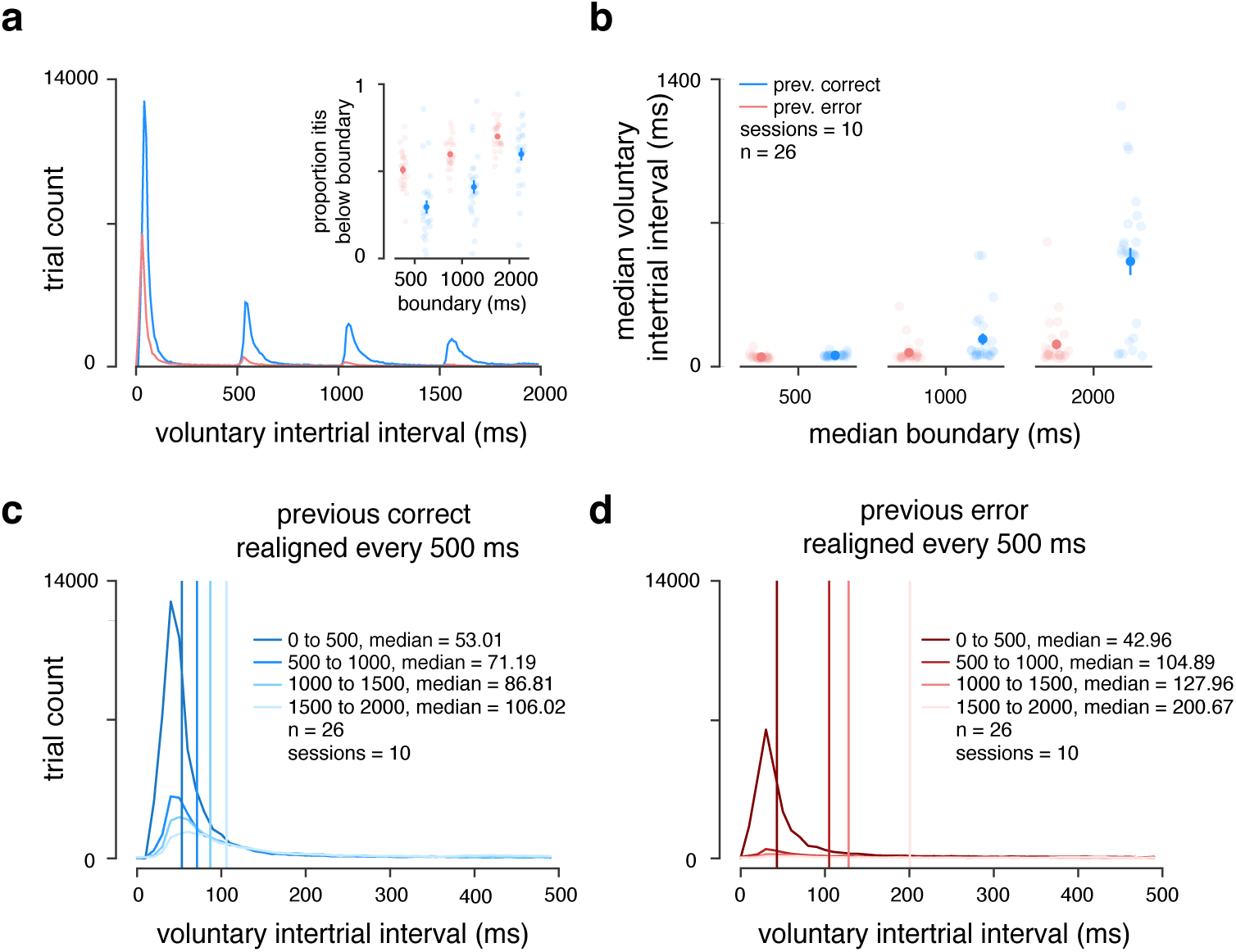
Analysis of voluntary intertrial intervals. **(c)** Histogram of voluntary ITIs (time in addition to mandatory experimentally determined D_*err*_ and D_*corr*_) for n = 26 rats across 10 sessions for previous correct (blue) and previous error (red) ITIs. Voluntary ITIs are spaced every 500 ms because of violations to the ‘cannot lick’ period. *Inset* : proportion of voluntary ITIs below 500, 1000 and 2000 ms boundaries. **(d)** Median voluntary ITIs up too 500, 1000 and 2000 ms boundaries. **(e)** Overlay of voluntary ITIs spaced 500 ms apart after previous correct trials. **(f)** Overlay of voluntary ITIs spaced 500 ms apart after previous error trials.

**Figure S13:**
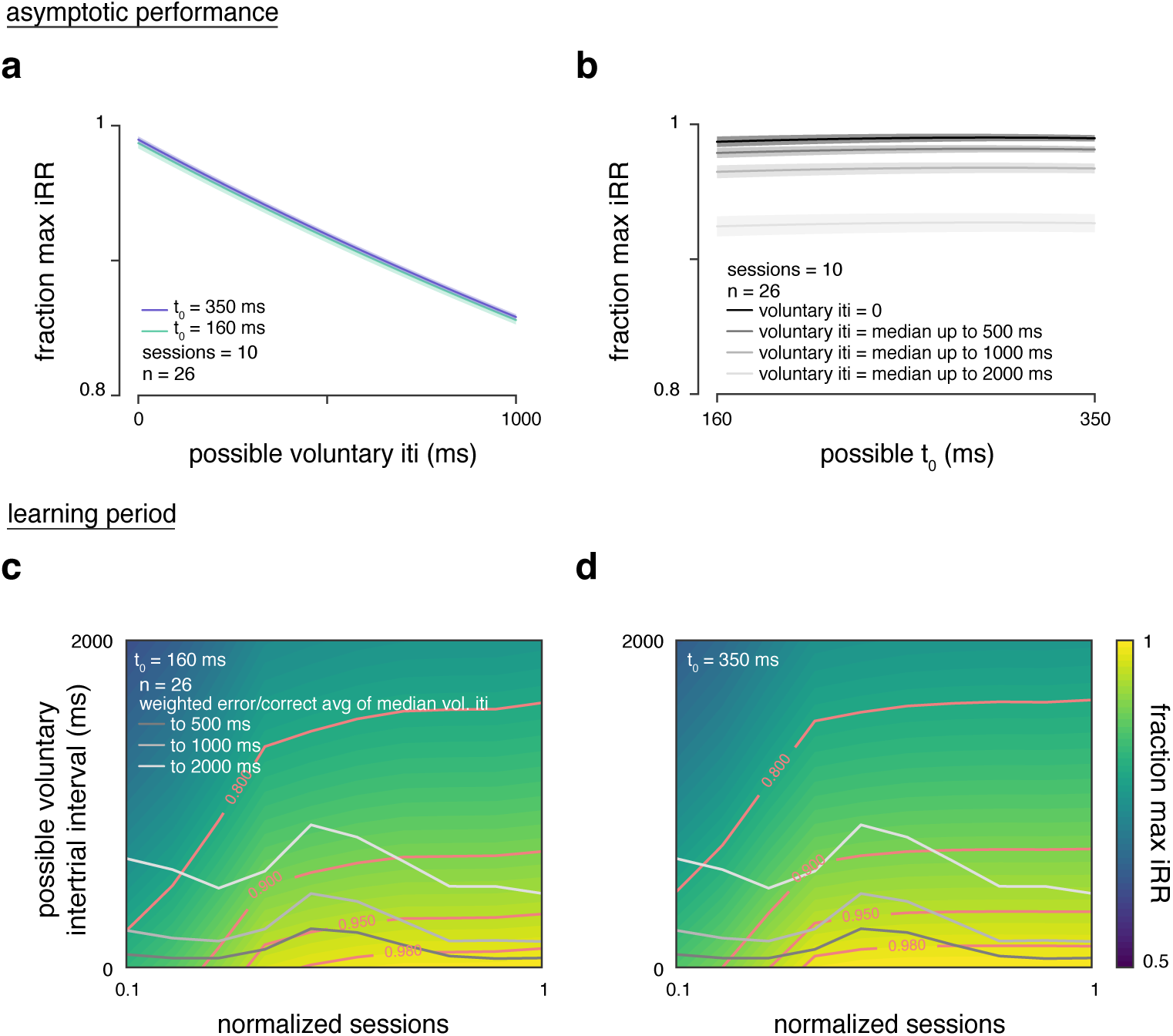
Reward rate sensitivity to T_0_ and voluntary inter-trial interval. **(a)** Fraction of maximum instantaneous reward rate across n = 26 rats over 10 sessions at asymptotic performance over possible voluntary ITI values of 0 - 1000 ms and over the minimum and maximum estimated t_0_ values. **(b)** Fraction of maximum instantaneous reward rate across n = 26 rats over 10 sessions at asymptotic performance over possible t_0_ values from 160 - 350 ms (min to max estimated t_0_ values) and over the median voluntary ITIs with 500, 1000 and 2000 ms boundaries. **(c)** Fraction of maximum instantaneous reward rate across n = 26 rats as a function of normalized training time during learning period and possible voluntary ITIs from 0 to 2000 ms calculated with the t_0_ minimum of 160 ms. The grey curves represent a weighted average over previous correct/error median voluntary ITIs over normalized training time. Contours with different fractions of maximum instantaneous reward rate in pink. **(d)** Same as in **c** but calculated with t_0_ maximum of 350 ms.

